# Epigenomic complexity of the human brain revealed by single-cell DNA methylomes and 3D genome structures

**DOI:** 10.1101/2022.11.30.518285

**Authors:** Wei Tian, Jingtian Zhou, Anna Bartlett, Qiurui Zeng, Hanqing Liu, Rosa G. Castanon, Mia Kenworthy, Jordan Altshul, Cynthia Valadon, Andrew Aldridge, Joseph R. Nery, Huaming Chen, Jiaying Xu, Nicholas D. Johnson, Jacinta Lucero, Julia K. Osteen, Nora Emerson, Jon Rink, Jasper Lee, Yang Li, Kimberly Siletti, Michelle Liem, Naomi Claffey, Caz O’Connor, Anna Marie Yanny, Julie Nyhus, Nick Dee, Tamara Casper, Nadiya Shapovalova, Daniel Hirschstein, Rebecca Hodge, Boaz P. Levi, C. Dirk Keene, Sten Linnarsson, Ed Lein, Bing Ren, M. Margarita Behrens, Joseph R. Ecker

## Abstract

Delineating the gene regulatory programs underlying complex cell types is fundamental for understanding brain functions in health and disease. Here, we comprehensively examine human brain cell epigenomes by probing DNA methylation and chromatin conformation at single-cell resolution in over 500,000 cells from 46 brain regions. We identified 188 cell types and characterized their molecular signatures. Integrative analyses revealed concordant changes in DNA methylation, chromatin accessibility, chromatin organization, and gene expression across cell types, cortical areas, and basal ganglia structures. With these resources, we developed scMCodes that reliably predict brain cell types using their methylation status at select genomic sites. This multimodal epigenomic brain cell atlas provides new insights into the complexity of cell type-specific gene regulation in the adult human brain.

---

The brain is the most complex organ in the human body consisting of billions of neuronal and non-neuronal cells, characterized by extensive diversity in gene expression, anatomy and functions. High throughput epigenomic profiling has been used to elucidate the gene regulatory programs underlying such cellular complexity (*1–3*), in both normal and disease brain states. Considered the fifth base of DNA, 5’-methylcytosines (5mCs) are the most common modified bases in mammalian genomes, providing an important epigenetic mechanism for regulating gene expression. Most 5mCs in vertebrate genomes occur at cytosine-guanine dinucleotides (CpGs). In vertebrate neuronal systems, however, 5mCs are also abundantly detected in non-CG (or CH, H=A, C, or T) contexts (*4*). Both CG- and CH-methylation (mCG and mCH, respectively) are highly dynamic during brain development and show remarkable cell type specificity (*1, 5, 6*). mCG and mCH are both essential for gene regulation and brain functions (*7*). In addition to DNA methylation, gene expression also requires proper conformation of chromatin folding (3D chromatin conformation), which is organized into active or repressive chromatin compartments, topologically associating domains (TADs) and chromatin loops (*8*). Such spatial organization facilitates the interaction between gene promoters and their regulatory elements (e.g., enhancers), providing additional critical layers of regulatory mechanisms. DNA methylation and chromatin conformation interplay and coordinate in regulating gene expression and these processes are highly correlated (*3*).

Here, we profiled both DNA methylation and chromatin conformation in a large number of adult human brain cells from cortical and subcortical regions using single-nucleus single and multiomic epigenomic sequencing technologies. These epigenomic cell maps complement single cell/nucleus transcriptome-based brain cell census approaches by providing a genome-wide view of DNA methylation and chromatin organization in distinct brain cell types, allowing an assessment of the cell-type specific function of noncoding regions in the human genome (*9, 10*). Moreover, such epigenomic information not only allows annotation of cell type-specific regulatory elements but also provides a comprehensive description of the unique and dynamic nature of 3D chromatin structures found in cell types across brain regions (*1*, *2*).

## Epigenome-based brain cell type taxonomies

We dissected 46 brain regions covering brain structures of the cerebral cortex (CX, 22 regions), basal forebrain (BF, 2), basal nuclei (BN, 11), hippocampus (HIP, 5), thalamus (THM, 2), midbrain (MB, 1), pons (PN, 1) and cerebellum (CB, 2) (Fig. 1A, Fig. S1A, and tables S1 and S2). Most regions had independent biological replicates from three adult male donors (table S3) except for two regions in the amygdala (BM and CEN; two donors for each) due to the dissection difficulty and tissue availability (Fig. S1A and table S1). Fluorescence-activated nuclei sorting (FANS) was used to isolate 90% NeuN-positive and 10% NeuN-negative cells in each sample to enrich neurons (Fig. S1A). We profiled genome-wide DNA methylation in each cell using snmC-seq3 (“mC”)(*11*) with samples from all 46 brain regions. Additionally, we also simultaneously profiled DNA methylation and chromatin conformation from individual cells using snm3C-seq3 (“m3C”)(*3*) with samples from 17 brain regions from CX, BF and BN (Fig 1B, and Fig. S1A). In total, 378,940 mC and 145,070 m3C nuclei passed rigorous quality control measures (Fig. S1B; see Methods). On average, 0.94 million stringently filtered reads were produced per mC cell and 2.20 million reads per m3C cell, covering 3.3% and 5.4% of the cytosines of the human genome, respectively. On average, 406,000 chromatin contacts were detected per m3C cell. These levels of genome coverage enabled reliable quantification of the DNA methylation levels of genomic features (Fig. S1C), identification of methylation variable regions, and determination of TADs and chromatin loops in distinct cell types throughout the brain.

**Figure 1.**
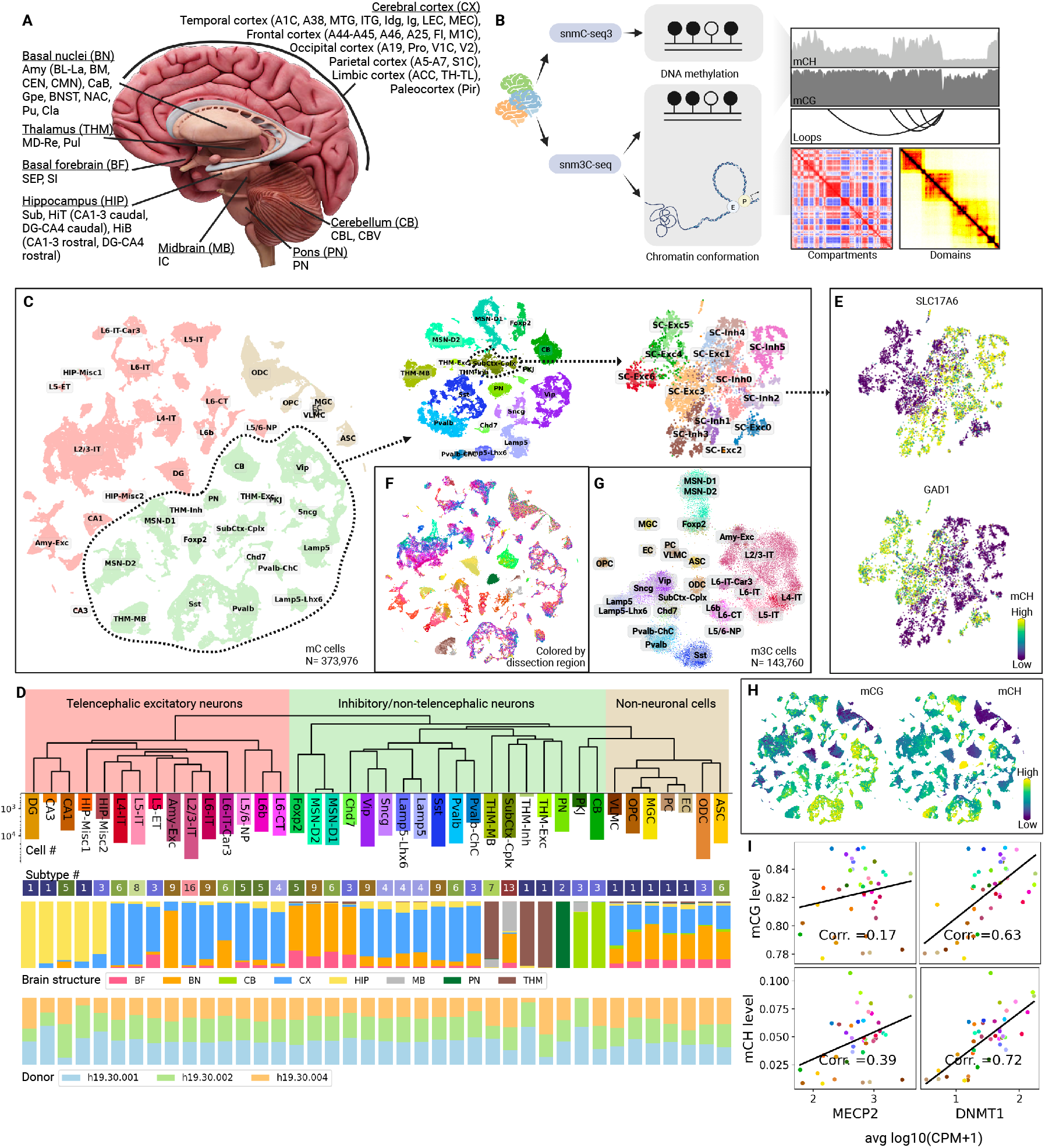
Epigenomic profiling of human brain cells with snmC-seq3 and snm3C-seq. (A) Human brain structures and regions covered in this study. (B) Schematics of profiling aspects of snmC-seq3 and snm3C-seq. (C) Iterative clustering and annotation of human brain nuclei. Cells from the whole mC dataset, from the inhibitory/non-telencephalic neuron cell class, and from the SubCtx-Cplx major type are visualized successively using t-distributed stochastic neighbor embedding (t-SNE), colored by the cell groups annotated in the corresponding iterations. (D) The robust dendrogram of the major types and the meta info of subtype numbers, brain structure, and donor origins. (E) The SubCtx-Cplx major type has both excitatory and inhibitory subtypes, marked by CH-hypomethylated genes SLC17A1 and GAD1, respectively. (F) Human brain cells are colored by the dissection regions. (G) 2D visualization of brain nuclei profiled by snm3C-seq. The visualization was generated from chromatin conformation data in 100kb resolution. (H) Global CG- and CH-methylation levels vary across brain cell types. (I) Global methylation levels of major types correlate with DNA methylation reader MECP2 and writer DNMT1.

We performed iterative clustering analysis on the mC dataset using CG- and CH-methylation levels on 100 kb bins throughout the human genome (Methods). The nuclei were first classified into three cell classes: telencephalic excitatory neurons, inhibitory/non-telencephalic neurons, and non-neuronal cells (Fig. 1, C and D). Each cell class was further divided into major cell types (referred to as “major types” through this paper) and annotated based on CH-hypomethylated gene markers for neuronal cell types and CG-hypomethylated markers for non-neuronal cell types. Where possible, cell types were annotated using the nomenclature for known brain cell types previously described in the literature (e.g. (*12*)); otherwise, cell clusters were annotated according to either the regional composition or distinct marker genes of the cell type. One caveat of the former approach is that cell types annotated using marker genes defined in rodents or non-human primates might not reflect the corresponding gene activity in human cell types. For example, the gene SNCG is lowly expressed in the human major type corresponding to mouse Sncg cells. Nevertheless, using common nomenclature aids in crossspecies comparison and existing knowledge transfer. We identified 40 major types from the three cell classes, which were further divided into 188 cell subtypes (Fig. 1D, Fig. S2, A to C, and tables S4 and S5). We performed subtype clustering using stringent parameters, requiring each subtype to have at least five unique marker genes compared with all other subtypes (Methods). Despite some cell proportion variation, all major types and subtypes were conserved across donors (Fig. 1D and Fig. S2C).

While non-neuronal major types have a relatively even distribution across brain structures, major neuronal types show considerable spatial specificity (Fig. 1, D to F). Most major telencephalic excitatory neurons are specifically grouped by spatial location. Excitatory neurons from the hippocampus are divided into separate clusters according to their origins in the sub-structures of the hippocampus (CA1, CA3, & DG). Cortical excitatory neurons cluster together according to both their cortical layers (L2/3, L4, L5, L5/6, & L6; L=layer) and projection types (IT = intratelencephalic, CT = corticothalamic, ET = extratelencephalic, and NP = near-projecting). The basal nuclei excitatory neurons mainly originate from the amygdala and form one specific group. The major type L6-IT-Car3 consists of neurons originating from the deep layers of the cortex and claustrum of the basal nuclei. Although they have similar DNA methylation patterns at the major type level, a recent study in mice showed that L6-IT-Car3 cells have divergent projection patterns (*13*). Telencephalic inhibitory neurons formed eleven major types, seven from mostly cortical regions (Pvalb, Pvalb-ChC, Sst, Lamp5, Lamp5-Lhx6, Sncg, and Vip) and four from basal nuclei and basal forebrain (MSN-D1, D2, Foxp2, and Chd7). In the thalamus, we identified one excitatory and two inhibitory major types. One inhibitory major type, THM-MB, shares similar DNA methylation profiles with a small population of cells in the midbrain. The other inhibitory major type, THM-Inh, is very rare; we detected only 361 cells in the entire dataset (0.07% of brain cells profiled). As THM-Inh features hypomethylation in genes EPHB1 and EPHA1 (Fig. S2D), this major type could originate from the habenular nuclei of the thalamus, possibly due to contamination during dissection. Neurons from Pons formed one specific major type (PN). The cerebellum contains two highly specific major types: Purkinje cells (PKJ), a rare cell type in the brain (867 cells, 0.17%), and cerebellar granular cells (CB). Some populations of cells from the basal nuclei and the midbrain form a conspicuous major type SubCtx-Cplx, which shows striking heterogeneity among its subtypes. For example, SubCtx-Cplx consists of both excitatory and inhibitory cells (Fig. 1E), while all other major neuronal types contain only one of them. SubCtx-Cplx also features highly variable DNA methylation among subtypes in the genes of neurotransmitter receptors, transporters, and neuropeptides (Fig. S2E).

The cell types determined from single nucleus DNA methylation profiles were corroborated using two orthogonal data types: single-cell transcriptome (scRNA-seq) data (see companion manuscript Siletti et al. (*14*)) and single-cell chromatin accessibility data (snATAC-seq; see companion manuscript Li et al.) produced using nuclei prepared from dissection regions from the same three human brains (Methods). Integrative analysis between these datasets reveals the strong correspondence between cell types determined using different molecular modalities (Fig. S3A). Notably, all epigenome-based cell subtypes correspond well with transcriptome-based clusters (Fig. S3B), though they were derived from ~10 times more cells and from ~2 times more brain regions. The DNA methylation-derived cell type labels were transferred to the matching scRNA and snATAC cells to facilitate further multi-omic joint analysis (Methods).

The robust dendrograms constructed from the single nuclei DNA methylation profiles show similarities between major types and subtypes (Fig. 1D and Fig. S2C; Methods). Telencephalic excitatory and inhibitory/non-telencephalic neurons are well-separated from non-neuronal cells, each type forming a specific clade (except CB and PKJ). CB and PKJ were grouped with the non-neuronal cell types in the dendrogram, likely owing to their similar global levels of CG- and CH-methylation (Fig. 1H and Fig. S4A).

Global methylation levels varied across major cell types ranging from 77.7% to 85.5% for CG-methylation and from 0.8% to 10.7% for CH-methylation. Non-neuronal and granular major types (DG and CB) have the lowest global levels in CG- and CH-methylation, consistent with the previous study in mice (*1*). Cortical inhibitory neurons have the highest CG-methylation levels, while several non-telencephalic neurons, particularly from the thalamus, midbrain, and pons, show the highest CH-methylation levels (Fig. 1H and Fig. S4A). Interestingly, cell-type global methylation levels correlate with the gene expression levels of DNA methylation readers and modifiers (Fig. 1I and Fig. S4B). The expression level of MECP2 and DNMT3A, the major CH-methylation reader and writer, show positive correlations (ρ=0.39, 0.35) with the global CH-methylation levels in contrast to a weaker correlation with CG-methylation (ρ=0.17, 0.08; Fig. 1I and Fig. S4B). The CG DNA methyltransferase gene DNMT1 showed a positive correlation (ρ=0.63) between its expression levels and global mCG across cell types (Fig. 1I), matching its role as the major mCG maintainer in mature neurons (*15*). On the other hand, we unexpectedly observed an even higher correlation between DNMT1 expression levels and mCH levels (ρ=0.72, Fig. 1I) compared with mCG; though DNMT1 is thought to have little effect on mCH (*16*). These findings suggest an unknown relationship between DNMT1 and mCH or some yet-to-be-discovered factor influencing both DNMT1 expression and mCH.

To obtain consistent cell type annotation for the cells profiled by m3C, we iteratively co-clustered the m3C nuclei with mC nuclei and transferred cell type labels from annotated mC nuclei to m3C nuclei (table S6; Methods). With the improved scHiCluster framework (*17*), we not only readily separated nearly all major cell types (exceptions being MSN-D1 and MSN-D2) using the chromatin contact only (Fig. 1G) but also revealed, for the first time, considerable regional diversity on the single-cell embedding (Fig. S2F). These results suggest considerable cell type and regional specificity of 3D chromatin structures.

## Chromatin organization in human brain cells

Chromatin is organized into structures at different scales. The subchromosomal-level compartment brings together regions that are tens to hundreds of megabases (Mb) away, while TADs and chromatin loops are driven by interactions within several megabases (*18*). To examine the cell-type specificity of genome folding at different lengths scales, we first analyzed the proportion of contacts detected at different genome distances within each single cell. Intriguingly, we observed that neurons are highly enriched for shorter-range interactions (<2Mb), while in most non-neural cells, longer-range contacts (>20Mb) are enriched (Fig. 2, A to C). Astrocyte and oligodendrocyte progenitor cells show intermediate interaction ranges, while mature oligodendrocytes are more similar to the non-neural cells (Fig. 2, A and B). In addition, cortical and subcortical excitatory neurons are more enriched for shorter-range interactions than cortical inhibitory cells (Fig. 2, A and B). The ratio between shorter and longer interactions is highly correlated with the number of RNA molecules detected in the cell type (ρ=0.87, Fig. S5A), indicating the contact distance could be associated with gene expression activity. These findings suggest that the contact distance spectrum, which is usually thought to be a signature of cell-cycle phases (*19*), can also be highly cell-type dependent.

**Figure 2.**
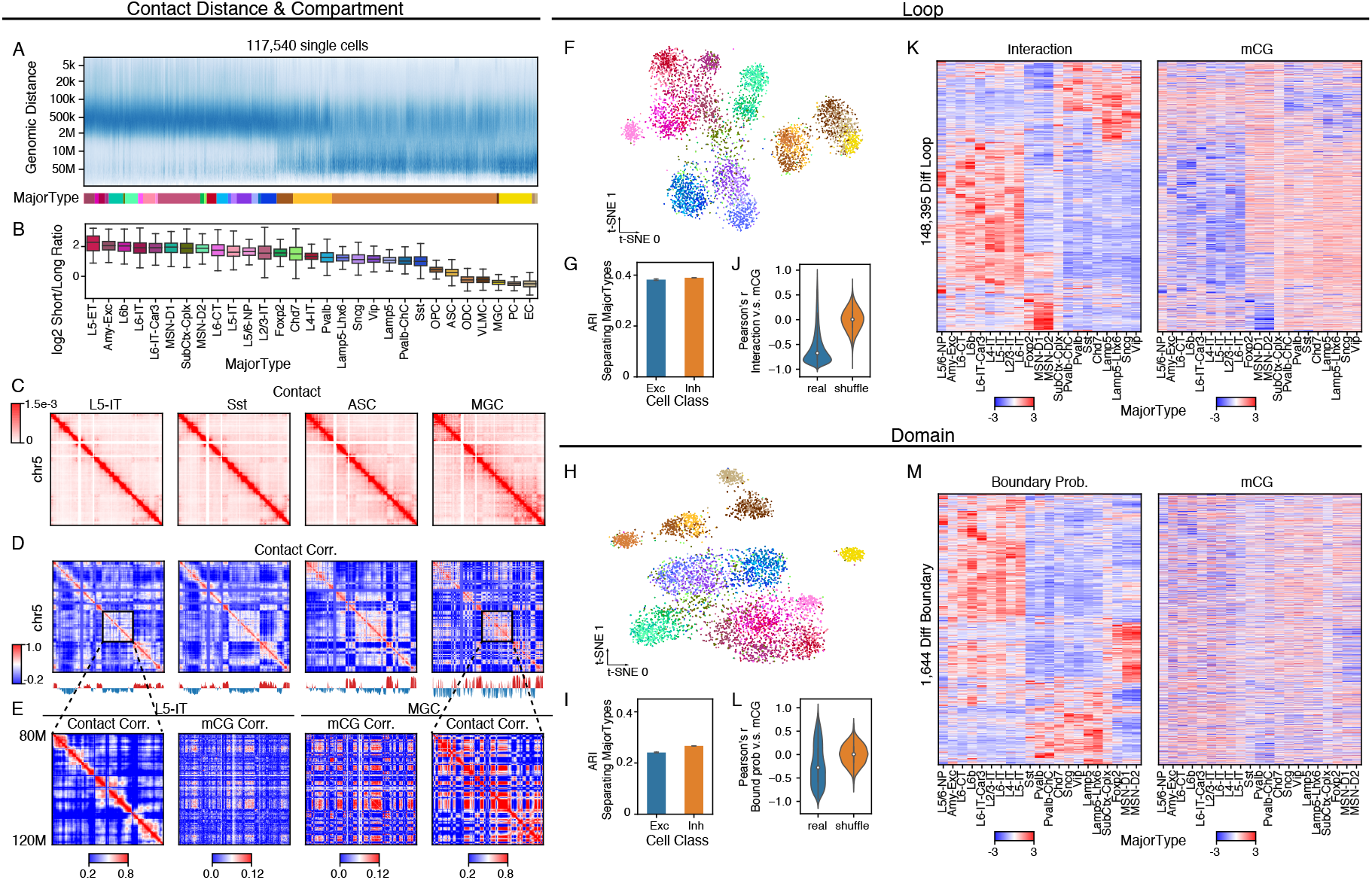
Diversity of 3D genome structures across major types. (A) Contact frequency against genomic distances across single cells. The bottom color bar shows which major type the cell belongs to. (B) Log2 ratio between shorter range and longer range contact frequency in each major type, ordered by the median ratio over cells. (C) The imputed contact maps of four major types. (D) The heatmaps show the correlation matrices of distance normalized contact maps in (C), and the line plots show the first principal component of the correlation matrices, with 0 as the middle line to separate red and blue. (E) Zoom in view of two matrices in (D) and the corresponding correlation matrices of CG methylation across cells. (F, H) t-SNE plot of cells (n=5,707) using loops (F) or domains (H) as features, colored by major types. (G, I) The adjusted rand index (ARI) between cortical excitatory major types or cortical inhibitory major types and clusters identified with loop (G) or domain (I) based cell embedding. Bars represent mean ± s.e.m of K-Means with 10 random seeds. (J) The imputed contact strength of differential loops and the average mCG level of the two loop anchors. (K) The distribution of Pearson’s correlation between two matrices in (J) across rows. (L) The boundary probability of differential domains and the mCG level at the boundaries. (M) The distribution of Pearson’s correlation between two matrices in (L) across rows. For all box plots, the center line denotes the median; box limits denote the first and third quartiles, and whiskers denote 1.5 × the interquartile range.

To investigate whether the increase in long-range chromatin interactions in non-neuronal cells leads to stronger compartmentalization, we identified chromosome compartments within each major type at 100 kb resolution (Fig. 2D). This analysis revealed that the enrichment of longer-range interactions showed little correlation with the compartment strength (Fig. S5, B and C), indicating that the enriched long-range interactions include both intra- and inter-compartment interactions. We next hypothesized that the methylation status of two genome loci is more likely to change together if they are physically closer and have stronger 3C interactions. Indeed, the co-methylation coefficient matrices (correlation of methylation levels between two genomic bins across single cells) form plaid patterns that resemble the compartment structures defined by chromatin contacts (Fig. 2E and Fig. S5D), suggesting the genome is segregated into local co-methylation domains as well as two sets of co-methylation regions across the chromosome. Notably, this coregulation structure was also observed with single-cell ATAC-seq data for chromatin accessibility (*20*), which, together with our data, provides independent evidence for genome compartmentalization. To further test the association between DNA methylation and 3D genome architecture, we compute the correlation between the chromatin interaction strengths and the average methylation levels of the interacting anchors across single cells at 100 kb resolution. We noticed that these correlations are also connected with chromosome compartments (Fig. S5E), where the negative correlations are more likely to be observed in the active compartment (A Compartment; p-values<1e-300, Wilcoxon Rank-Sum test; Fig. S5F).

Next, we identify domains in single cells using the imputed contact matrices at 25 kb resolution. More domains are identified in neurons (median 4,813) than in non-neurons (median 4,308, p-values<1e-300, Wilcoxon Rank-Sum test) (Fig. S5G). We defined the boundary probability of a genomic bin as the proportion of cells in which the bin is determined as a domain boundary. The recurrent domain boundaries across single cells correspond to the domain boundaries identified with cell-type pseudo-bulk contact maps (Fig. S5H). We also identified chromatin loops at 10 kb resolution in each of the 119 cell subtypes with >=100 m3C cells (median 541,551 loop pixels, with 59,905 loop summits). In total, 24.3% of loops represent interactions between distal DMRs and annotated gene promoters (TSS±2kbp), 38.1% are distal DMR-DMR loops, and 5.8% are promoter-promoter loops (Fig. S5I). Using either domain boundary probabilities or loop strengths was able to recapitulate the hierarchical structure of cell type similarities (Fig. S5, J and K) and separate single cells by cell types in the embedding spaces (Methods; Fig. 2, F and H), suggesting the cell type specificity of these 3D structural features. Chromatin domains showed relatively weaker power to distinguish finer scale major types within excitatory or inhibitory cell class (Fig. 2, G and I). This finding indicates that different scales of 3D features may play a role in the gene regulation specification at different granularities of cell types, where the DNA loops we identified are more major type-specific than domains.

To investigate the cell-type-specific 3D structural features more closely, we identified 1,644 differential domain boundaries across the major cell types. Chromatin domains had been considered conserved across cell types (*21–24*).However, in agreement with more recent studies showing domain dynamics across cell types and development (*3*, *25–27*), our data suggest that domains may differ even between similar cell types (Fig. S5H). We also identified 148,395 differential loops across major types. Promoter-DMR loops show higher cell-type specificity than promoter-promoter or DMR-DMR loops (Fig. S5L). The strength of loops is strongly anticorrelated with DNA CG methylation (Fig. 2, J and K), and this anticorrelation tends to be stronger at loops with higher variability across cell types (Fig. S5M). The boundary probability is also anticorrelated with CG methylation; however, the correlation is weaker compared with loops (Fig. 2, L and M). The anticorrelation could have resulted from the effect of DNA methylation on the binding of factors driving genome folding (e.g., CTCF) or the recruitment or exclusion of methylation writer/eraser (e.g., DNMT, TET) through loop formation. Further developmental or mechanistic studies are needed to resolve a causality relationship and its possible interplay with gene expression. Together, these analyses reveal the cell-type specificity of chromatin architecture and its relationship with DNA methylation at an unprecedented cell-type resolution in the human brain.

## Cell-type specific DNA methylation patterns and associated gene regulatory landscapes

To delineate the cell type-specific methylation profiles, we identified 24,455 CH- and 13,096 CG-pairwise differentially methylated genes (DMGs) among the subtypes (Fig. S6A; Methods). We also identified 2,059,466 CG-differentially methylated regions (DMRs) in the human genome from 188 brain cell subtypes (Fig. 3A; Methods). These cell typespecific DMG and DMR patterns depict distinct DNA methylation signatures for brain cell identities. In addition, they provide critical information to understand gene regulatory programs in brain cells. Both CG- and CH-methylation levels of DMGs negatively correlate with their gene expression levels (*5*, *28*, *29*), while DMRs predict candidate cis-regulatory elements (cCREs) (*6*, *28*)), and motifs enriched in their sequence indicate the cell typespecific transcription factors (TFs; (*29*)).

**Figure 3.**
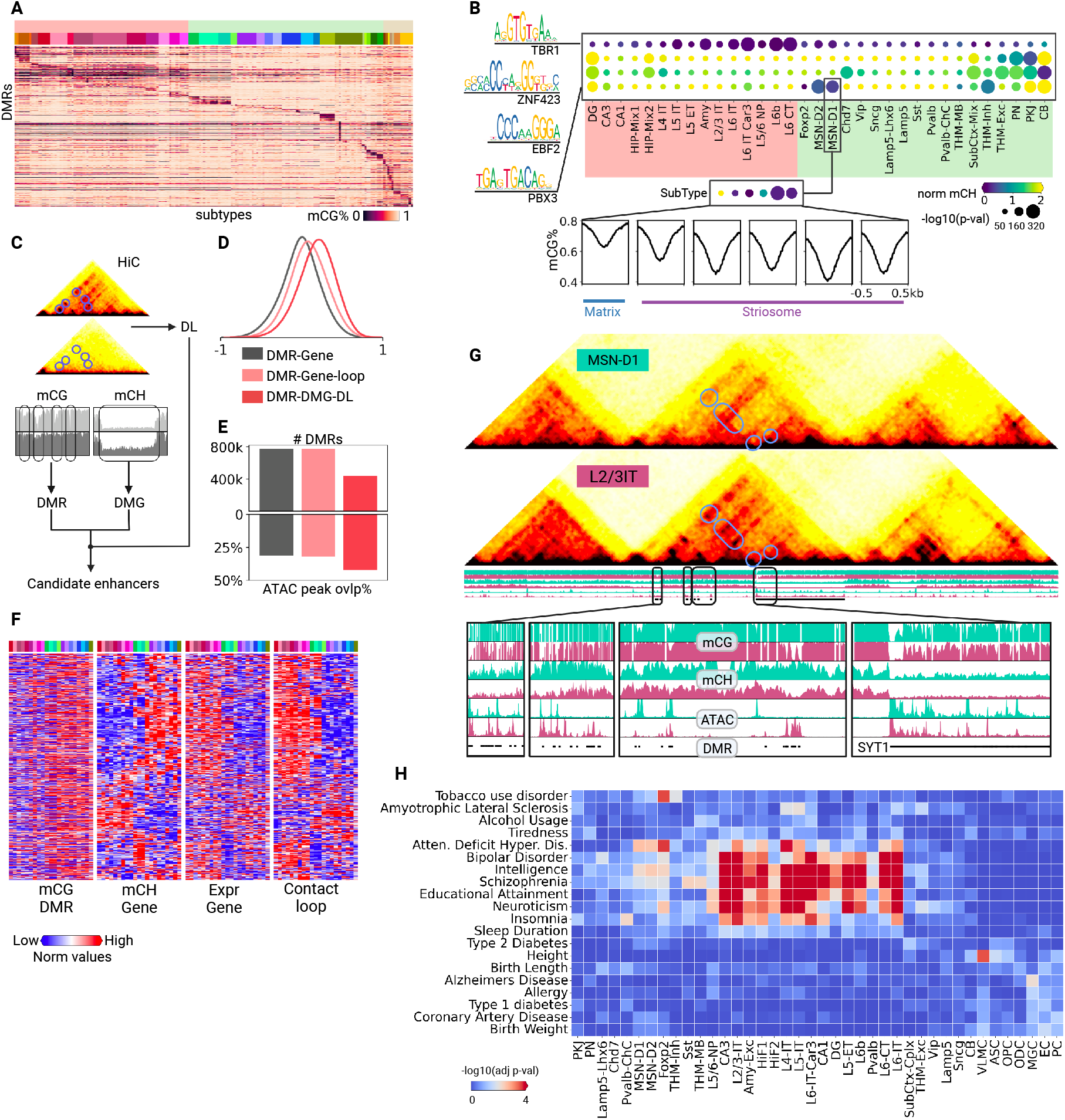
Gene regulation in brain cells. (A) Methylation levels of cell type-specific DMRs across 188 cell subtypes. (B) CH-hypomethylated transcription factors and the enrichment of their motifs in CG-hypomethylated DMRs. The lower panel shows average methylation levels of transcription factor PBX3 in its potential binding sites across the whole genome. (C) Workflow of determining candidate enhancers. (D) The putative enhancers predicted with the integration of DMRs, DMGs, and differential loops have a higher correlation with the corresponding genes in methylation. (E) Open chromatin regions are enriched in the putative enhancers. (F) The candidate enhancers and the target genes (anti-) correlated with each other in their methylation, expression, and contact strength. (G) The gene SYT1 in major types L2/3-IT and MSN-D1 vary in gene body CH-methylation, DMR CG-methylation, and chromatin organization in 3D space. (I) Heatmap showing the results of linkage disequilibrium score regression analysis of the variants associated with the indicated traits or diseases in DMRs identified from major human cell types.

To determine putative TFs in each cell type, we performed a motif enrichment analysis for cell type-specific hypomethylated DMRs (hypo-DMRs; see Methods). Combining with the DMG analysis results, we assigned TFs to specific cell types if a TF has its gene body hypomethylated in the cell type and its binding motif enriched in hypo-DMRs of the cell type (Fig. S6B; Methods). In total, 612 TFs were assigned to major neuronal types and subtypes, where they potentially play important roles in shaping and maintaining cell identities. For example, the TF TBR1 which plays a fate-determining role in the development of corticofugal projection neurons (*30*) was assigned to deep-layer excitatory neurons, particularly L6-CT and L6b (Fig. 2B). The TFs ZNF423 and EBF2 were assigned to Purkinje cells. ZNF423 was additionally assigned to cerebellar granular cells (Fig. 2B). Both TFs are crucial in the proper development of the cerebellum, and EBF2 particularly directs the migration of Purkinje cells (*31–33*).

In addition to major types, analysis of subtypes can further provide insights into subtle differences in TF utilization among subtypes. For instance, the TF PBX3, assigned to the MSN-D1 major type, is only hypomethylated in the subtypes showing hypo-methylation at MEIS2 and FOXP1 genes, two marker genes of striatal striosome compartment (Fig. S6C). This suggests that PBX3 may also be expressed only in the striosome but not in the matrix compartment (Fig. 3B). This finding agrees with the previous observation that PBX3 is preferentially expressed in striosomes, and could be crucial for striatum development (*34*, *35*). Furthermore, examination of the genome-wide potential binding sites (overlapped regions between hypo-DMRs and TF motifs) of PBX3 also shows that the average methylation levels are lower in striosome subtypes compared to the matrix (Fig. 3B), suggesting a spatial-specific regulating role of this TF in the striatum.

We further integrated information on DMGs, DMRs, and different loops to identify putative enhancers in each cell type (Fig. 3C). DMRs were associated with genes if the TSS of the gene was within 5 Mb of the DMR. Pearson correlations between mCG levels of DMRs and mCH levels of genes body across cell subtypes were calculated to assess the association. The associations are slightly stronger between DMRs and DMGs than between DMRs and other genes (Fig. S6D). We further filtered the DMR-DMG pairs and kept only the ones that overlapped with both anchors of a loop or a differential loop (DL). Both loop- and DL-filtered DMR-DMG pairs show stronger associations (Fig. 3D and Fig. S6D). DL-filtered DMRs have the strongest association with paired genes (Fig. 3D) and an increasing overlapping ratio with open chromatin regions (Fig. 3E), indicating their putative regulating roles in adult brain cells. Putative regulating DMR/gene pairs (3.2M) were identified between 1,122,919 DMRs and 12,327 genes. These DMRs, their paired genes, and the corresponding loops were (anti-) correlated (Fig. 3F) and together may collectively orchestrate cell type-specific gene regulatory programs. For example, the gene SYT1 encodes Synaptotagmin-1, an integral membrane protein of synaptic vesicles, which serves as the major switch allowing neurons to release neurotransmitters at synapses when sensing calcium ions (*36*).Distal DMRs hypomethylated in L2/3-IT are connected to the promoter region of SYT1 through DNA loops which are either depleted or weakened in MSN-D1 cells. Concordant with such regulation differences, SYT1 has a higher expression level and lower gene body mCH level in L2/3-IT than MSN-D1 (Fig. 3G and Fig. S6E). Thus, integration of CG- and CH-methylation, as well as the chromatin conformation, help delineate specific cell type regulatory programs.

Thousands of non-coding loci have been identified as associated with brain diseases via genome-wide association studies (GWAS), many of which reside in enhancer regions (*37*). DMRs and loops can position genetic variants in regulatory elements of specific cell types. Using linkage disequilibrium score regression (*38*), we found significant associations between 20 brain diseases or complex traits (Methods) and DMRs or loop-overlapping DMRs in human brain cells (Fig. 3H and Fig. S6F; Methods). Risk variants related to schizophrenia, bipolar disorder, and neuroticism show strong enrichment in hypomethylated DMRs of excitatory neurons located in the cortices and hippocampus. Alzheimer’s disease (AD) is associated with microglia (MGC; Fig. 3H; (*39*)). Risk variants of tobacco usage disorder are enriched in a dopamine neuron cell type Foxp2 from basal ganglia (Fig. 3H), a brain region related to tobacco addiction (*40*).

## Regional heterogeneity in cortices and basal ganglia

In addition to the distinct cell type specificity for the brain anatomical structures discussed above, we also observed that cells of the same type from different regions within one brain structure also manifest considerable DNA methylation diversity. Prominent functional and gene expression heterogeneity across the cortex has been described in previous studies, particularly in the neocortex and hippocampus. Gradients of morphogens in the developing mouse neocortex play crucial roles in early cortical region specialization (*41*). Microarray profiling showed that the transcriptomes of two neocortical regions are more similar if the two regions are closer spatially (*42*). Recent single-cell studies also revealed that the regional heterogeneity is cell type dependent in the mouse brain, which was particularly prominent in neocortical excitatory neurons in gene expression and gene body DNA methylation (*1*, *43*). In addition to the neocortex, our dataset encompasses a broader range of brain regions, including the allocortex (e.g. Pir) and periallocortex (e.g. LEC and MEC), allowing investigation of the gene regulatory elements underlying the transcriptome diversity across these brain regions.

To distinguish regional diversity from other confounding sources of heterogeneity within one cell type, we developed a workflow to extract the regional landscape hidden within single-nuclei DNA methylation profiles (Fig. 4A). By combining single nuclei DNA methylation profiles and the corresponding brain region information, we transformed the cells from the DNA methylation space to a “joint methylation-brain region space” (Fig. 4A; Methods), representing from which brain regions cells sharing similar methylation profiles were dissected. Specifically, if two cells are closer in this space, their methylation neighbors (neighboring cells in the methylation space) are more likely to come from similar brain regions. Therefore, a cell’s trajectory in this space represents a transition across brain regions accompanied with changes in their methylomes. By emphasizing the regional methylation similarity rather than simply capturing the major axes of methylation variation, the transformed embedding of single cells facilitates further analyses examining regional effects on DNA methylation in brain cells.

**Figure 4.**
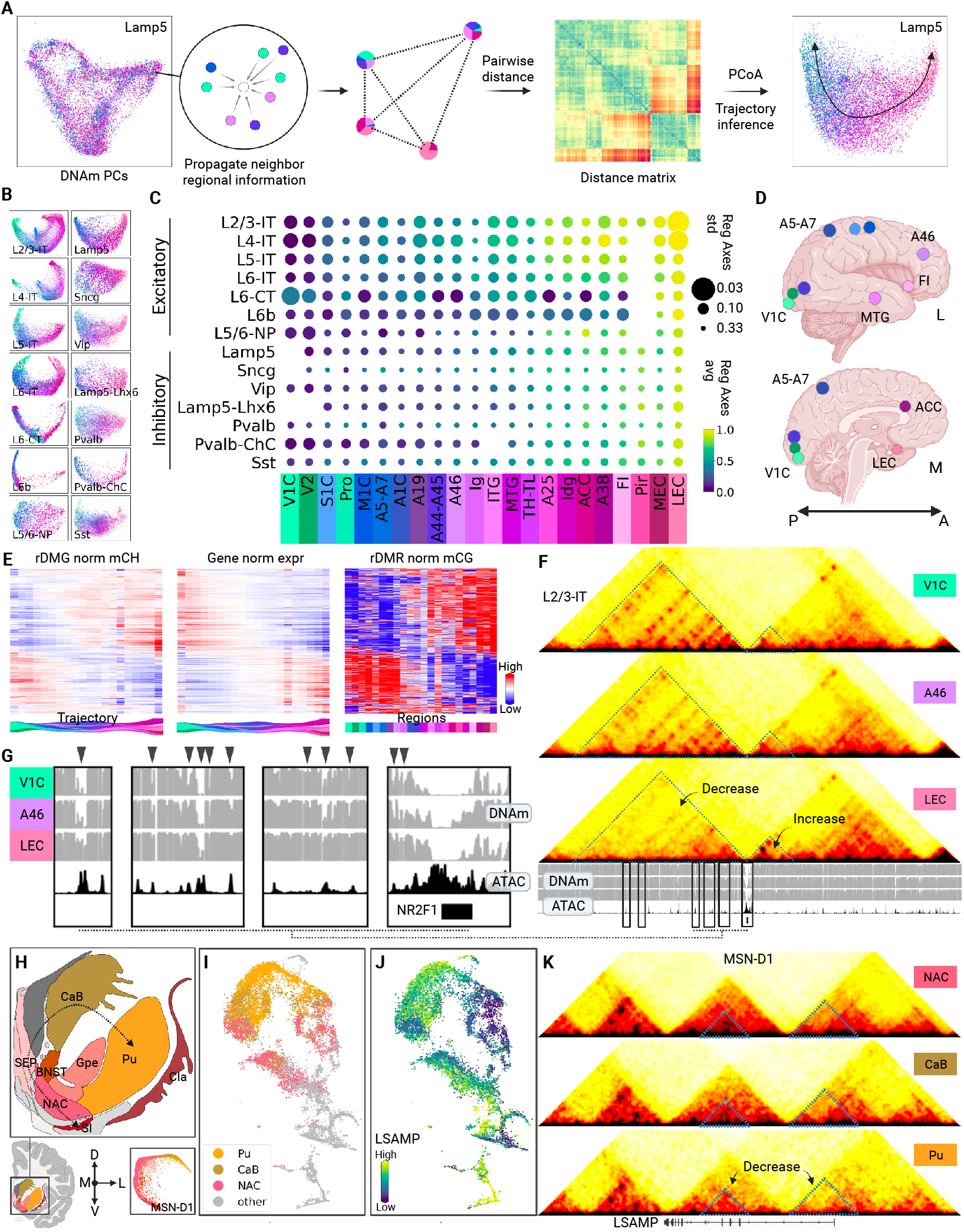
Regional axes of cortical and subcortical cells. (A) Workflow of determining regional axis from single nucleus DNA methylome. (B) 2D visualization of cortical neurons in regional spaces. The cortical neurons were colored by their dissection locations. (C) Trajectory analysis shows a common axis among cortical neurons. The scatter plot shows how regional indices vary in each cortical region. (D) Schematic of example cortical dissection locations. (E) Example of rDMGs in L2/3-IT cells. The rDMGs show gradient methylation levels along the axis we detected, which (anti-) correlate with the gene expression and the methylation levels of nearby rDMRs. (F) Chromatin conformation around the gene NR2F1 shows gradient changes in domain and loop strength. The two associated chromatin domains change in opposite directions. (G) Zoom-in view of example differentialloop-overlapping rDMRs from the “decreasing” domain. The methylation levels increase from V1C to A46 to LEC. (H) Inhibitory neurons in basal ganglia show an L–D–V axis in DNA methylation (I) 2D t-SNE visualization of MSN-D1. Cells from NAC, CaB and Pu are highlighted. (J) Gene body methylation of LSAMP in MSN-D1 shows the difference among cells from NAC, CaB and Pu. (K) Chromatin conformation around the gene LSAMP in MSN-D1. Cells from NAC and CaB show stronger chromatin interactions in LSAMP compared to Pu.

Cortical excitatory neurons showed remarkable regional diversity in their DNA methylation profiles, especially the intratelencephalic-projecting neurons (LX-IT; Fig. 4B). The regional diversity of cortical inhibitory neurons (*44*) was less studied due to their inconspicuous regional patterns in transcriptome and epigenome (*1*, *43*, *45*). Our analysis uncovered their regional diversity: despite being less profound, cortical inhibitory neurons showed considerable regional differences (Fig. 4B). The embeddings in the regional spaces showed a shared gradient among cortical neuronal cell types, both excitatory and inhibitory (Fig. 4B). We further derived regional axes for each cell type in the aforementioned joint methylation-brain region spaces using a single-cell trajectory analysis tool STREAM (*46*) to quantitatively analyze the gradient molecular diversity across regions. Although the regional axes may have certain discrepancies among cell types, they followed a common Posterior–Lateroanterior–Medioanterior (P–LA–MA) trend: from the posterior part of the brain (e.g., the primary visual cortex V1C) to the anterior lateral part (e.g., the prefrontal cortex A46 & the middle temporal gyrus MTG) and then to the anterior medial part (e.g., the anterior cingulate cortex ACC & the lateral entorhinal cortex LEC; Fig. 4, C and D). Only one cell type, L6-CT, shows an exceptional regional pattern (Fig. 4, B and C) from the P–LA–MA axis. Nevertheless, the shared pattern allows us to merge the axes of different cell types into a consensus regional axis for cortical neurons (Fig. 4C; Methods).

Epigenetic features vary along the regional axis, which could involve the regional specification of brain cortices. For example, the transcription factor NR2F1, also known as COUP-TFI, is a crucial morphogen in brain development, which establishes the caudal-rostral neocortical regional specialization (*41*) and the boundary between the neocortex and the entorhinal cortex (*47*). NR2F1 has relatively high expression levels in V1C (P) and LEC (MA) while low in A46 (LA; Fig. S7A). The gene body CG- and CH-methylation levels show a negatively correlated pattern with its expression (Fig. 4G and Fig. S7B). Interestingly, NR2F1 is associated with two chromatin domains whose strengths change in opposite directions from V1C to A46 to LEC (Fig. 4F). The upstream domain shows stronger interactions in V1C, and DMRs within this domain are hypo-methylated in V1C compared to LEC. Contrarily, the downstream gene domain shows stronger interaction and harbors DMRs hypo-methylated in LEC (Fig. 4G, and Fig. S7, C and D). These coherent regional variations in epigenetic features and transcription activity imply a regulatory domain switching for NR2F1 from V1C to A46 to LEC, which needs further investigation. Systematic examination of the regionally differential epigenetic features determined in total 14,606 (average 2.9k within each major type) from cortical neuronal cell types, 885.4k (63.2k) regional DMRs (rDMRs; Fig. S7G; Methods), 773k (71.2k) regional differential loops and 1,495 (136) regional differential domains (Fig. S7E). Many rDMGs and rDMRs show monotonic methylation gradients along the P–LA–MA axis (Fig. 4E, and Fig. S7, F, and G), while more complex patterns (such as NR2F1) also exist. These multiomic analyses provide an epigenetic basis for the regional diversity of gene expression in the brain cortex and allow the identification of key enhancers and regulatory programs in the specialization of cortical regions.

The inhibitory neurons in basal ganglia showed remarkable regional diversity as well. Four major cell types were identified in the regions of human basal ganglia, including D1- and D2-type medium spiny neurons (MSN-D1 and MSN-D2), and two cell types named Foxp2 and Chd7, featuring CH-hypomethylation of the genes FOXP2 and CHD7 respectively. Our analysis reveals a profound molecular regional axis from lateral to dorsal to ventral parts (L–D–V) of the basal ganglia (Fig. 4H). We also found regional differences in gene regulation along this regional axis of basal ganglia. For example, the gene of Limbic System Associated Membrane Protein (LSAMP) shows a methylation gradient: CH methylation levels elevated from NAC to CaB to Pu (Fig. 4, I and J), accompanied by the decrease of strengths of chromatin domains and loops around this gene (Fig. 4K). In total, we determined 6,371 (2.7k) rDMGs and 398.8k (99.7k) rDMRs from all four cell types (Fig. S7E). Two cell types, MSN-D1 and MSN-D2, had enough m3C cells for regional differential analysis on chromatin conformation, in which we identified 98,276 (50,271) rDLs and 193 (99) rDDs (Fig. S7E). Most rDMGs and rDMRs (anti-) correlate strongly with the L–D–V axis (Fig. S7, H and I), indicating that regional diversity is a critical source of within-cell-type heterogeneity in the basal ganglia. As the major component of the basal ganglia, the striatum comprises the ventral part, including the nucleus accumbens (NAC), and the dorsal part, including caudate (CaB) and putamen (Pu). The two parts were known to differ in both functions and neural connections (*48*, *49*). Our data and analysis provide the epigenetic basis of the D-V difference. In addition, we extended the D–V axis and revealed regional differences in a finer resolution within the dorsal part of the striatum (Fig. 4H).

## Conservation of brain cell types and DMRs between humans and mice

Overall conservation of cortical cell types between primates and rodents was noted in comparative single-cell transcriptomic studies of several cortical regions (*50*, *51*). Whether the conservation of taxonomies holds in broader brain regions remains less studied. To address this question, we performed an integration analysis comparing the single nuclei DNA methylation profiles between human and mouse brains (*1*). Corresponding brain regions between the two species were used, including the cerebral cortex, basal forebrain, basal nuclei, and hippocampus (Methods). Twodimensional embedding of the integration shows a general matching between cell types of the two species (Fig. 5A and Fig. S8A). However, three major types defined from human brains are noticeably discrepant with mouse brain cells. Mouse L4-IT neurons aligned only to subpopulations of their human counterparts (Fig. 5B), suggesting a larger heterogeneity in human L4-IT neurons compared to mice and confirming findings from a previous comparative study between human MTG and mouse V1 (*50*). The human hippocampal major type HIP-Misc1 was integrated with a population of mouse cortical IT neurons instead of hippocampal neurons. Another human hippocampal major type HIP-Misc2 did not match any mouse brain cell type. Integration analysis against the parallel scRNA study (*14*) validates these two human hippocampal major types, which both have the signatures of DNA hypomethylation and RNA expression of the transcription factor TSHZ2 and its overlapped long non-coding RNA (lncRNA) AL109930.1 (Fig. 5C and Fig. S8B). Although two unmatched hippocampal major types will need further investigation, taxonomies at the major type level are generally conserved across broader brain regions between humans and mice. These matched cell types show consistently higher global levels in both CG- and CH-methylation in humans than in mice (Fig. 5D and Fig. S8C).

**Figure 5.**
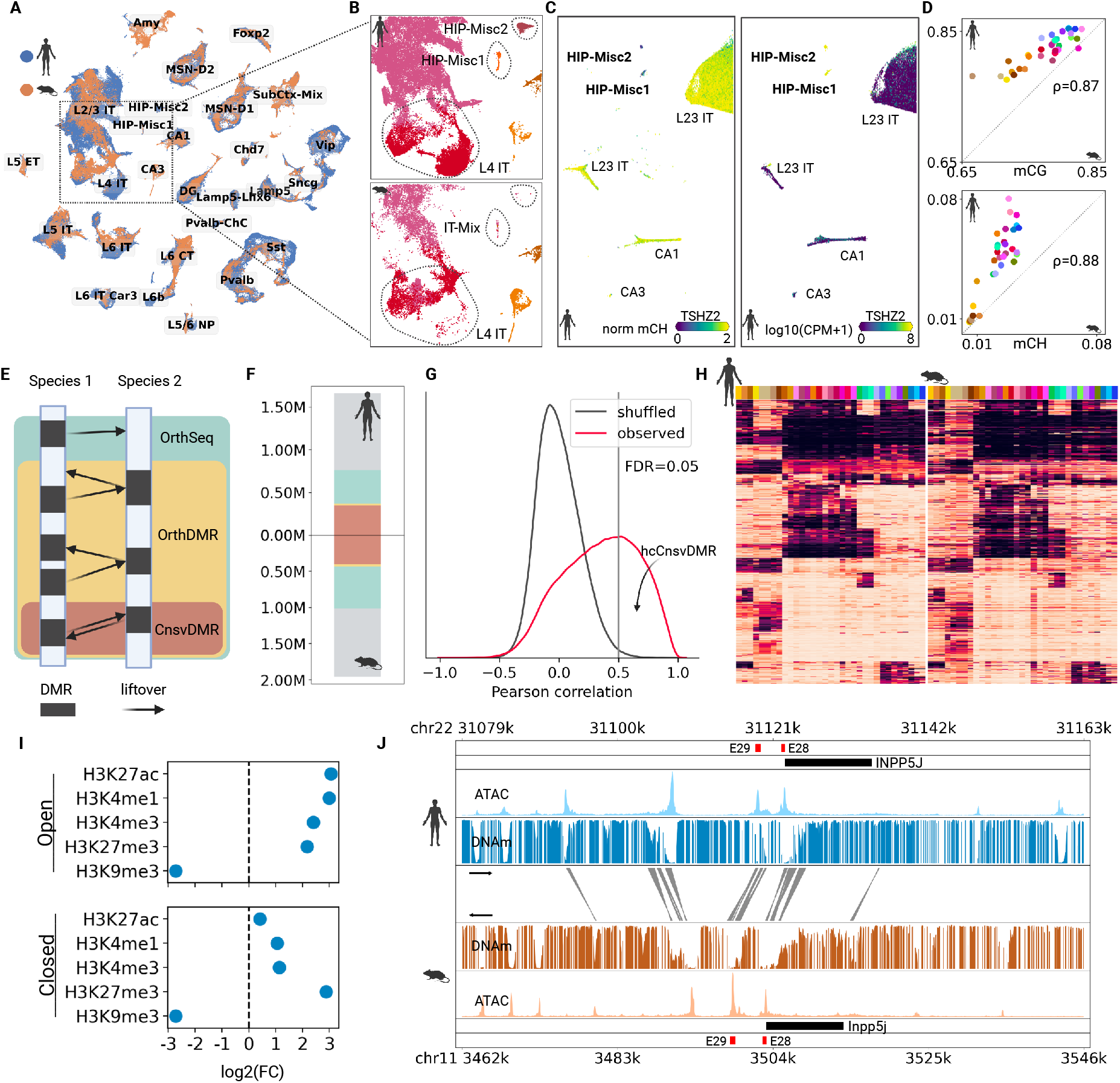
Cross-species comparison between human and mouse brain cell methylomes. (A) Integration of single-cell methylomes between human and mouse brains shows general cell type conservation across species. Neuronal cell types are visualized using 2D t-SNE. (B) Discrepancy between cell types of human and mouse brains in cell types L4-IT, HIP-Misc1, and HIP-Misc2. (C) Integration between human brain DNA methylation and transcriptomic datasets validates the cell types of HIP-Misc1 and HIP-Misc2, which both feature CH-hypomethylation and gene expression of TF TSHZ2. (D) Conserved cell types between human and mouse show correlated global CH- and CG-methylation levels. Methylation levels in human cells are higher than in mouse cells. (E) Schematic of cross-species matching of cell type DMRs. (F) Overall, ~50% of DMRs have orthologous sequences in the other species, among which ~25% are reciprocal DMRs. (G) Distribution of cross-species correlation of DMR methylations. DMR methylation in humans and in mice is highly correlated compared to the randomly shuffled background. (H) Examples of methylation levels of hcCnsvDMRs. (I) The hcCnsvDMRs show enrichment in the histone modification marks. The open hcCnsvDMRs are likely to be active enhancers and promoters while the closed ones are likely to be vestigial enhancers. (H) Browser view of hcCnsvDMRs around gene INPP5J in major type Pvalb. The regions colored by red are the cell type-specific distal enhancers validated in Ref (*29*).

To compare the gene regulation between human and mouse brain cells, we first identified major type-specific hypo-DMRs within each species and used liftOver to match DMRs in corresponding major types between species (Fig. 5E). Then, we defined three hierarchical DMR categories based on direct and reciprocal matching: 1) OrthSeqs, which have ortholog sequences in the other species; 2) OrthDMRs, the OrthSeq whose ortholog sequences are also hypomethylated DMRs in the other species; and 3) CnsvDMRs, the orthDMRs that can be liftOvered reciprocally between the two species (Fig. 5E). In general, 40~60% DMRs are in the category of OrthSeqs across cell types, around half of which are OrthDMRs (Fig. 5F and Fig. S8D), and most (95%) of OrthDMRs are also CnsvDMRs (Fig. 5F and Fig. S8D). Furthermore, the methylation levels of CnsvDMRs showed a significant correlation between humans and mice across cell types (Fig. 5, G and H). We further compared the observed correlations against the cell-type-wise randomly shuffled background to select CnsvDMRs with significant correlation across species, which we called hcCnsvDMRs (Fig. 5G). Functional enrichment analysis of hcCnsvDMRs shows that they are highly enriched in biological processes related to forebrain development and in cellular components related to dendrites and synapses (Figs. S8, F and G; Methods). Comparison to histone modifications in mouse forebrains (*6*) demonstrated these DMRs are depleted from heterochromatic regions (H3K9me3) as well as enriched in regions of enhancers (H3K27ac & H3K4me1), promoters (H3K4me3), and poised enhancers (H3K27m3; Fig. S8E; Methods). Categorizing the DMRs further into open or closed status based on their chromatin accessibility (*2*) shows that open DMRs are enriched in enhancers and promoters. In contrast, closed DMRs are particularly enriched in the poised enhancers (Fig. 5I), which were probably active during development.

The high conservation of DMR methylation levels between species points to a strategy of enhancer discovery through comparative epigenetics. For example, INPP5J, a gene expressed specifically in Pvalb cell type, has a number of distal and proximal hypomethylated hcCnsvDMRs, which overlap with matched chromatin-accessible regions as well (Fig. 5J). Two of these CREs were previously validated as specific enhancers that can be used in viral tools to target and manipulate Pvalb cells (Fig. 5J) (*52*).

## Single-cell methylation barcodes (scMCodes) reliably predict human brain cell type identification

Cytosine methylation variation in the genomes of cells contains molecular “engrams” in the form of hyper- and hypo-methylated regions that may represent past and present gene regulatory events (*53*). Methylation variation at singlenucleotide resolution has successfully been utilized for a wide array of applications, such as predicting epigenetic ages (*54*), tracing cell lineage (*55*, *56*), and diagnosing lifethreatening diseases like cancer from circulating free DNA (or cell-free DNA, cfDNA) (*57*, *58*). We have shown that DNA methylation levels of gene bodies or 100kb genome bins can effectively distinguish brain cell types (*1*). In addition to these relatively long features, we observed many CpG sites with methylation patterns highly specific to certain cell types in the human brain. For example, the CpG site at chr1:46203766 is fully methylated in all cells other than THM-MB, while the CpG site at chr18:36273621 is unmethylated in all DG cells (Fig. S9B). With such high specificity at both single-cell and single CpG site levels, we derived single cell Methylation barCodes (scMCodes) for human brains, which determine brain cell types using the methylation states of a number of CpG sites at single-cell resolution (Fig. 6A and Fig. S9A; Methods). We iteratively selected cell type-specific and high-confidence CpG sites that distinguish brain cell types (Methods). The CpG sites were further clustered into 39k groups according to their across-cell-type methylation patterns and filtered by their cell-type predicting power through a Random Forest model (Fig. S9A; Methods). We selected 800 groups with a total of 12k CpG sites as the scMCode (Fig. 6B, and tables S7 and S8) to achieve high enough cell type predicting power (Fig. 6C) while maintaining a relatively small feature number (Fig. S9C). This scMCode achieved ~93% cell type prediction accuracy (Fig. 6D; Methods). To ensure the scMCode approach is robust, we conducted cross-donor tests by deriving scMCodes and training the predicting model using methylation data from one donor while testing cell type predictions using the others. These cross-donor predictions achieved high accuracies (92~93%) among all training-test pairs (Fig. 6E), demonstrating that the scMCode method is a robust and reliable predictor of cell type identity across individuals. Because single-cell sequencing has limited genomic coverage, not all CpG sites are expected to be covered in each single cell. Indeed, on average only ~200 CpG sites from different feature groups are actually used in each cell to achieve such high prediction accuracy (Fig. 6F), indicating the effectiveness of scMCode in determining human brain cell types from a few hundred select methylation sites.

**Figure 6.**
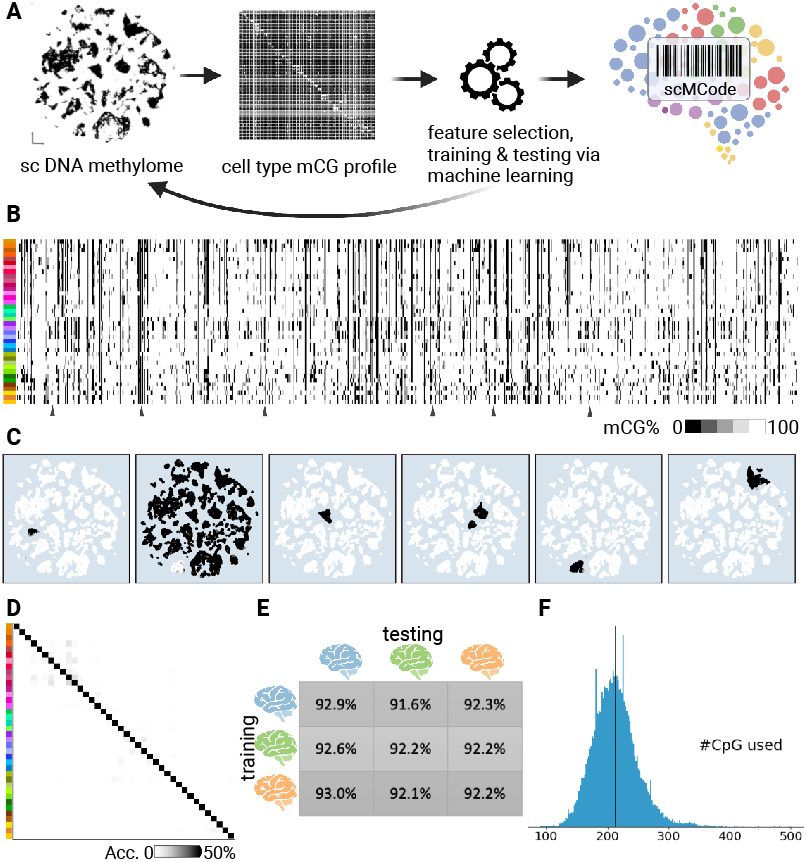
snMCodes for brain cell types. (A) Workflow of deriving snMCodes. (B) snMCodes derived from all three donors. (C) The methylation status of snMCode features is highly specific to cell types. (D) Heatmap of confusion matrix shows snMCodes can accurately predict cell types. (E) snMCodes are robust to individual differences in determining brain cell types. (F) snMCodes predict human cell types with a limited number of CpG sites at single-cell resolution.

## Discussion

Knowledge of cell type diversity in the human brain and their specific gene regulatory programs is essential for understanding brain functions and developing therapeutics for brain disorders. Here we report comprehensive profiling of DNA methylation and chromatin conformation at singlecell resolution in the human brain, comprising 524,010 deep coverage nuclei from 46 cortical and subcortical brain regions. This dataset allows the identification of 188 epigenetically distinct cell types and the characterization of their molecular features, including marker genes, differentially methylated regions, signature transcription factors and DNA loops. Furthermore, integrative DNA methylation and chromatin conformation analysis can effectively distinguish the cell-type-specific candidate regulatory elements and their interaction with target genes. Moreover, the diversity of 3D genome structures was revealed at different scales and with unprecedented cell type resolution.

Taking advantage of the broad dissection coverage in brain sampling and in-depth epigenome sequencing coverage for each nucleus, we could delineate regional axes in both cortex and basal ganglia, which provides a novel approach for investigating brain region heterogeneity. Cross-species comparison between human and mouse brain cell methylomes reveals conservation in cell types and regulatory elements, suggesting an enhancer discovery strategy via comparative epigenomics.

The rich and precise regulatory information stored in cytosine methylation allowed the derivation of scMCodes for human brain cells, providing a novel approach for reliable cell type identification using the methylation status of a small number of select CpG sites. Analysis of circulating-free DNA (cfDNA) methylation represents a non-invasive method for cancer diagnosis (*58*). cfDNA methylation has also been implicated as promising biomarkers for brain disorders (*59*). Our scMCode approach could be directly applied in profiling and analyzing cfDNA, which provides potential means for brain disorder diagnosis, pathological region identification, and treatment selection.

Overall, this multimodal human brain cell atlas complements transcriptomic studies of cell type diversity, regional heterogeneity, and evolution conservation with an underlying epigenetic basis, providing both a valuable resource for the neuroscience community to study gene regulation in brain cells and the raw material (enhancers) to develop new genetic tools for targeting specific cell types.

## Acknowledgments

We thank Jesse R Dixon, Trygve E Bakken, Nikolas L Jorstad, and Song-Lin Ding for the scientific discussion and suggestive comments. We are very grateful to members of the Ecker group for feedback and discussion. We thank Dr. Erica Melief and Aimee Schantz for outstanding administrative assistance, Lisa Keene, Katelyn Kern, and Amanda Keen for support in tissue collection, and most importantly the brain tissue donors and their loved ones, without whom this work would be impossible.

## Author contributions

Study supervision: J.R.E. Contribution to data generation: W.T., J.Z., A.B, R.G.C., M. K., J.A., A.A., J.R.N., H.C., N.D.J., J.L., J.K.O., A.P., N. E., J.R., J.L., M.L., N.C., C.O., C.V., A.M.Y., J.N., N.D., T.C., N.S., D.H., R.H., B.P.L., C.D.K. Contribution to data analysis: W.T., J.Z. Q.Z., J.X. Contribution to data interpretation: W.T., J.Z., K.S., E.L., B.R., M.M.B., J.R.E. Contribution to writing the manuscript: W.T, J.Z., J.R.E. All authors edited and approved the manuscript.

## Data availability

The data analyzed in this study can be downloaded from http://neomorph.salk.edu/wtian/hba-data/. The data were produced through the Brain Initiative Cell Census Network (BICCN:RRID:SCR_015820) and deposited in the NEMO Archive (RRID:SCR_002001) under identifier nemo:dat-jx4eu3g accessible at https://assets.nemoarchive.org/dat-jx4eu3g. Raw and processed data were also deposited to NCBI GEO/SRA with accession number GSE215353. A browser of major type DNA methylome can be found here http://neomorph.salk.edu/hba/hba-majortype.php.

## Supplementary materials

Materials and Methods

Figs. S1 to S9

Tables S1 to S8

