## Supplementary material for "Epigenomic complexity of the human brain revealed by single-cell DNA methylomes and 3D genome structures": Methods

**This PDF file includes:**

Materials and Methods
Figs. S1 to S9
Tables S1 to S8

**Materials and Methods**:

**Human postmortem tissue specimens**

De-identified adult postmortem human brain tissue was obtained after receiving permission from the deceased's next of kin. Tissue collection was performed per the United States Uniform Anatomical Gift Act of 2006, described in the California Health and Safety Code section 7150 (effective 1/1/2008) and other applicable state and federal laws and regulations. In addition, the Western Institutional Review Board reviewed tissue collection procedures and determined that they did not constitute human subjects research requiring institutional review board (IRB) review.

Male donors 18–68 years of age with no known history of neuropsychiatric or neurological conditions were considered for inclusion in the study. Routine serological screening for infectious diseases (HIV, Hepatitis B, and Hepatitis C) was conducted using donor blood samples, and donors testing positive for infectious diseases were excluded from the study. Specimens were screened for RNA quality, and samples with average RNA integrity (RIN) values ≥7.0 were considered for inclusion in the study. Postmortem brain specimens were processed as previously described (*50*) (dx.doi.org/10.17504/protocols.io.bf4ajqse). Briefly, coronal brain slabs were cut at 1 cm intervals, photographed, frozen in dry-ice cooled isopentane, and transferred to vacuum-sealed bags for storage at -80°C until the time of further use. For the dissection of brain regions of interest, photos of tissue slabs were annotated by a neuroanatomist to outline regions to target for dissections. Then, tissue slabs were removed from the -80°C freezer and briefly transferred to -20°C, where they were held for ~1-3 hours to allow tissues to equilibrate to -20°C. Tissues were then transferred to a custom temperature-controlled cold table held at -20°C and the region of interest was removed using standard razor blades or scalpels. Tissue blocks were stored at -80°C in vacuum-sealed bags until later use.

**Nuclei isolation and Fluorescence Activated Nuclei Sorting (FANS)**

Nucleus isolation was conducted using a standard protocol as previously described (dx.doi.org/10.17504/protocols.io.y6rfzd6). Gating on DAPI and NeuN fluorescence intensity was as described previously (*50*). NeuN+ and NeuN- nuclei were sorted into separate tubes and were pooled at a defined ratio of 90% NeuN+ and 10% NeuN- nuclei after sorting. Sorted samples were centrifuged, frozen in a solution of 1X PBS, 1% BSA, 10% DMSO, and 0.5% RNAsin Plus RNase inhibitor (Promega, N2611), and stored at -80°C until further processing. The presorted nuclei pellets were defrosted and resuspended in DPBS+1%BSA, centrifuged, resuspended back in 1ml of DPBS, and sorted into 384-well plates. Nuclei from donors H19.30.001 and H19.30.002 were prepared and sorted into 384-well plates. For donor H19.30.004, frozen tissue blocks received from AIBS were processed following procedures previously described (*5*). Nuclei were labeled for NeuN fluorescence and sorted into 384-well plates as described (*1*).

**Library preparation and Illumina sequencing**

**snmC-seq library preparation.** snmC-seq3 libraries were prepared using an updated version of snmC-seq2. In brief, samples underwent bisulfite conversion and were barcoded with random primers. Samples were then pooled through two SPRI cleanups to compress 16 x 384-well plates into 1 x 96-well plates. Pooled samples were then adapted and amplified as previously described. Next, libraries were pooled and cleaned through two more SPRI cleanups. Finally, library concentrations were determined by Qubit and normalized for sequencing. snmC-seq3 and snm3C-seq (see below) libraries generated from human brain tissues were sequenced using an Illumina Novaseq 6000 instrument with S4 flowcells and 150 bp paired-end mode.

**snm3C-seq library preparation.** For some samples from donors H19.30.001 and H19.30.002, presorted nuclei were used. The presorted nuclei pellets were defrosted and resuspended in DPBS+1%BSA, centrifuged, and resuspended back in 1 ml of DPBS. For the remaining samples of donors H19.30.001 and H19.30.002 and all samples from donor H19.30.004, frozen tissue was pulverized using a mortar and pestle. All samples were then immediately crosslinked with 2% formaldehyde in solution for 5 min, quenched with 0.2M Glycine for 5 min, centrifuged and washed with DPBS, and stored at -80°C until ready for further processing. Next, nuclei were conditioned and digested using an Arima kit adapted for snm3C-seq for 1hr at 37°C, and 20 min at 65°C to inactivate enzymes, then ligated for 15min at room temperature. Finally, nuclei were resuspended in 1ml of DPBS+1%BSA, filtered through a 0.2 2μM filter, and sorted similarly to the snmC-seq3 samples.

**Donor-specific genomes**.

**gDNA Library prep protocol.** Genomic DNA was extracted from ground, frozen tissue using the DNeasy Blood and Tissue Kit (Qiagen, Valencia, CA). One μg of DNA was fragmented with a Covaris S2 (Covaris, Woburn, MA) to 300 bp, followed by end repair (Lucigen) and the addition of a 3' A base (New England Biolabs). Cytosine-methylated adapters provided by Illumina (Illumina, SanDiego, CA) we ligated to the sonicated DNA at 16˚C for 16 hours with T4 DNA ligase (New England Biolabs). Adapter-ligated DNA was isolated by two rounds of purification with AMPure X P beads (Beckman Coulter Genomics, Danvers, MA). The adapter-ligated DNA molecules were enriched by 4 cycles of PCR with the following reaction composition: 25μL of Kapa HiFi Hotstart (KapaBiosystems, Woburn, MA) and 5μl TruSeq PCR Primer Mix (Illumina) (50μlfinal). The thermocycling parameters were: 95˚C 2min, 98˚C 30sec, then 4cycles of 98˚C 15 sec, 60˚C 30 sec, and 72˚C 1min, ending with one 72˚C 5 min step. The reaction products were purified using AMPure X P beads. The purified PCR reactions of the adapter-ligation resulted in a library used for subsequent sequencing in Novaseq 6000.

**Variant calling from donor genome sequencing.** Whole genome sequencing reads were first QCed with the software fastp (v0.20.1) (*60*). The command line used is "fastp -i input_PE_R1.fastq.gz -I input_PE_R2.fastq.gz -o output_PE_R1.fastq.gz -O output_PE_R2.fastq.gz -w 4". The QCed reads were then mapped to human genome assembly GRCh38 (hg38) via the software BWA (v0.7.17)(*61*)with the

BWA-MEM algorithm with mapping results stored in bam format through the software samtools (v1.10)(*62*). Specifically, the command line used for mapping is "bwa mem -t 20 hg38-ref input_PE_R1.fastq.gz input_PE_R2.fastq.gz | samtools view -Sb - > output.bam".

The mapped reads were analyzed with the germline short variant discovery workflow of the Genome Analysis Toolkit (GATK, v4.1.8.1)(*63*). Briefly, we first removed the duplicated reads from the mapped reads, which then went through a base quality score recalibration step (BQSR) to generate analysis-ready reads. The variant references used in the BQSR step were dbSNP138, Mills and 1000 Genomes gold standard indels, and 1000 Genomes phase 1 SNPs. Next, candidate variants (SNPs+InDels) were called with the HaplotypeCaller of GATK from the analysis-ready reads and further filtered with a variant quality score recalibration step (VQSR) to determine the high-confidence SNPs and InDels, respectively. The variant references used to recalibrate SNP quality scores were Hapmap 3.3, OMNI 2.5, 1000 Genomes phase 1 and dbSNP138, and of InDels were Mills and 1000 Genomes gold standard indels and dbSNP138. All the references used in the BQSR and VQSR were downloaded from the GATK resource bundle (ftp://ftp.broadinstitute.org/bundle/hg38).

**Donor-specific reference genome.** For each donor, we selected the high-confidence homozygous SNPs using the function SelectVariants of GATK, and created donor-specific reference genomes by substituting the homozygous SNPs into the hg38 FASTA file using the function FastaAlternateReferenceMaker of GATK

**Common homozygous SNPs.** By comparing the homozygous SNPs of donors, we constructed a list of common SNPs shared among the three donors.

#### Mapping and count/feature matrix generation

For sequence read mapping of both snmC-seq3 and snm3C-seq datasets, we used our own custom pipeline (https://github.com/lhqing/cemba_data, version 1.2.1.dev94+gc65e173). The main steps of this pipeline included: 1) Demultiplexing FASTQ files into single-cell; 2) Reads level QC; 3) Mapping; 4) BAM file processing and QC; 5) final molecular profile generation. The details of the five steps were previously described (*11*). We mapped all of the reads to the donor-specific genomes. After mapping, we calculated the methylcytosine counts and total cytosine counts for two sets of genomic features in each cell. Non-overlapping chromosome 100kb bins of the hg38 genome (generated by "bedtools makewindows -w 100000"), were used for clustering analysis, and the genes defined by the human GENCODE v33 were used for cluster annotation and integration with datasets. Both CG and CH methylation levels of the features were normalized as previously described (*1*). The cell-by-feature matrices were generated from normalized methylation levels of each feature set.

#### Quality control measures

The sequenced cells were filtered based on these metrics: 1) mCCC% < 0.06; 2) global mCG% > 0.5; 3) global mCH% < 0.15; 4) total final reads > 250,000; 5) mapping rate > 0.5. For cells profiled with snm3C-seq, we required a cell to have > 50,000 cis contacts with a distance over 2500bp.

#### Clustering and annotation of snmC-seq3 data

**Clustering analysis.** CG- and CH-methylation levels of 100kb genomic bins were used as input features for clustering. We performed clustering analysis iteratively using the software package ALLCools (https://github.com/lhqing/ALLCools). In each iteration, the 100kb bins were first filtered by removing bins with mean total cytosine base calls < 250 or > 3000. Those who overlap with the ENCODE blacklist (*64*) were also excluded from the clustering analysis. The Top 5,000 highly variable features (HVFs) were then selected separately from both CG- and CH-methylation via support vector regression (SVR). We then applied principle component analysis (PCA) to each 5,000 features to reduce dimension. The top *n* principle components (PCs) were selected for each methylation type until there is no significant difference between the distributions of *n*-th and *(n+1)*-th PCs by two-sample Kolmogorov-Smirnov test with the criteria as the adjusted p-values < 0.1. We further performed pre-clustering for each top PC set and selected the PCs that are enriched in pre-clusters following Ref (*65*). Finally, the selected PCs from both CG and CH methylation PCs were concatenated for further analysis. We used Harmony (*66*) on the selected PCs in order to eliminate individual differences. The Harmonized features were further fed into the consensus clustering procedures previously described (*1*).

**Doublet/debris identification.** The read number of each cell in one plate is stable in both snmC-seq3 and snm3C-seq. Therefore we adopted a doublet/debris detection strategy based on cell relative reads to its plate. We first normalized the read number per cell to the mean reads of its plate. The cells with plate-relative-read numbers>1.2 or <0.8 were considered doublet/debris candidates. After each iteration of clustering, clusters would be labeled as doublet/debris if the cluster contained over 80% doublet/debris candidates and were eliminated from further analysis.

**Cell type annotation.** The clusters were manually annotated as major or subtypes according to their hypomethylated genes, which were either canonical brain cell type markers or determined *de novo* from the current dataset. We required each cell type to have at least five differentially methylated genes in CG and CH methylation compared to the other cell types. Otherwise, it would be merged with the closest cluster. A candidate cell type would be labeled as an outlier if all its cells were from a single donor.

**Clustering and annotation of snm3C-seq data**

To annotate cells from snm3C-seq, we combined them with the annotated snmC-seq3 cells and carried out an iterative clustering analysis similar to what was described above. The only difference was that batch effects from both individuals and sequencing technologies were corrected using the software Scanorama (*67*). After each clustering iteration, the cell type annotations were transferred from snmC-seq3 cells to snm3C-seq cells with a K Nearest Neighbor (KNN) classifier.

**Robust dendrogram of cell types**

We resampled a certain number of cells from each cell type without replacement to compute the average methylome profile for the cell type with genome features of 100kb-bins of both CG- and CH-methylation. The resampling number is 800 for major types and 500 for subtypes. The average profiles were then used to compute the pairwise correlation distances. This process was repeated 500 times to compute an average pairwise distance matrix, which was then used to construct the final cell-type dendrogram via hierarchical clustering with average linkage.

**Determine differentially methylated genes**

We determined the DMGs pairwise between cell subtypes for CG- and CH-methylation separately. To avoid potential bias caused by an imbalance of cell numbers of cell types, we downsampled cells in each cell subtype to no more than 500. All the protein-coding and long non-coding RNA genes (lncRNAs) defined by the human GENCODE v33 were tested for significant methylation decrease (or hypomethylation) using the Wilcoxon Rank Sum Test. The p-values were adjusted with multitest correction using the Benjamini-Hochberg procedure. We computed the Area Under the Receiver Operating Characteristic curve (AUROC) for the candidate genes. The genes with adjusted p-values≤0.001 and AUROC≥0.8 were considered pairwise DMGs in CG- and CH- methylation.

**Determine differentially methylated regions**

We merged single-cell DNA methylation profiles into the cell type (major type/subtype) profiles according to their cluster annotation in both donor-aggregated and donor-separated ways. Non-common homozygous SNP CpG sites of these methylation profiles were filtered out before further analysis. We then used the DMRfind function of the software MethylPy (v1.4.2; (*68*) ) to determine the mCG DMRs across all cell types of the donor-aggregated profiles. The command line used is "methylpy DMRfind --output-prefix OUTPUT_FILE_NAME --samples SAMPLE_NAMES --mc-type CGN --dmr-max-dist 250 --sig-cutoff

0.01 --allc-files MC_FILES". We further merged the successive DMRs if their distance is within 250bp and the Pearson correlation of their mCG fractions across 188 subtypes is greater than 0.8. We further screened each DMR by evaluating the reproducibility of the methylation pattern across cell types between donor-aggregated and -separated profiles. The evaluation criteria were 1) the Pearson's correlation coefficient between the mCG fractions across cell types is ≥0.5, and 2) the mean-absolute-error (MAE) is ≤0.1.

Each reproducible DMR was then assigned as hypo- or hyper-DMRs in each cell type based on the difference of its mCG fraction from its robust mean. The robust-mean m of each DMR was calculated by averaging the mCG fractions between 25th and 75 percentiles across cell types. The DMRs with mCG fractions greater than m+0.3 were assigned as the hyper-DMRs in each cell type, and lower than m-0.3 were assigned as hypo-DMRs. DMRs without any hypo- or hyper-DMR assignment were excluded from further analyses.

**Motif enrichment analysis**

746 transcription factor binding profiles (motif) from JASPAR2020 (*69*) were used to perform the motif enrichment analysis. Cell-type-specific hypo-DMRs were first segmented into 500bp bins, and then annotated with each motif by intersecting with the genome locations of the motifs. Motif genome locations were downloaded from http://expdata.cmmt.ubc.ca/JASPAR/downloads/UCSC_tracks/2020/hg38. To test for motif enrichment in the major cell types, hypo-DMRs were used as foreground signals, and the hypo-DMRs of all other major types were used as background. For testing at the cell subtype level, hypo-DMRs in only the other subtypes that belong to the same major type were used as background. The one-sided Fisher exact test was used to calculate the p-values of enrichment of the foreground against the background.

**Single-cell embedding based on chromatin contacts**

Single-cell contact matrices at 100kb resolution were imputed by scHiCluster (*17*) with pad = 1. The imputed contacts with distance >=100kb and <=1 Mb are used as features for singular value decomposition (SVD) dimension reduction. The first 30 principal components were normalized by singular values and L2 norms per cell, and then used for t-SNE visualization in Fig. 1G and S2F. To better visualize the heterogeneity of neuronal cells, we downsample each of the neuronal cell populations to 1,000 cells and, together with all the neurons to fit the t-SNE, and project back the other non-neuronal cells (*70*).

**Contact distance distribution analysis**

We generated a histogram of contacts for each single cell based on the distance between the two anchors of the contact. The bins are equally divided on the log2 distance scale, with a step size of 0.125, ranging from 2500 bp to 249 Mb (length of the longest chromosome). The i-th bin is the number of contacts with a distance between $2500 \times 2^{0.125i}$ and $2500 \times 2^{0.125(i+1)}$. In Fig. 2B and S5A, the short-long ratio was defined as the proportion of contacts in 51st (200k) to 76th (2M) bins divided by the proportion of contacts in 103rd (20M) to 114th (50M) bins.

**Compartment Analysis**

Pseudo-bulk contact matrices of each chromosome at 100kb resolution are used for compartment analysis. We first filter out the 100kb bins with abnormal coverage. Specifically, the coverage of bin i on chromosome c (denoted as $R_{c,i}$) is defined as the sum of the i-th row of the contact matrix of chromosome c. We only keep the bins with coverage between the 99th percentile of $R_c$ and twice the median of $R_c$ minus the 99th percentile of $R_c$. Contact matrices are normalized by distance, and Pearson's correlation matrices of the normalized matrices are used (*71*). We use the first principal component (PC1) of the correlation matrix and assign the compartment with higher CpG density (A compartment) positive scores to compute compartment scores for imputed matrices. For raw matrices, we use either PC1 or PC2, depending on which has a stronger correlation with CpG density because PC1 often corresponds to the chromosome arms. The results from the imputed matrices generally agree with the imputed matrices. However, the imputed matrices work better with smaller cell populations, while the raw matrices provide higher resolution when enough cells are merged. Therefore we use imputed matrices for individual cell type analyses and use raw matrices only for Fig. S5, D, and E to call compartments after merging all cell types. Saddle plots and compartment strengths are computed in the same way as described in (*72*). The average values of distance normalized contact matrices are used in Fig. S5B, and the average values of Pearson correlation between mCG levels of two bins across cells are shown in Fig. S5C.

**Identification of chromatin loops and differential loops**

Chromatin loops were identified with scHiCluster (*17*) in each major type, subtype, and major type within each brain region, respectively. To identify loops from a group of cells, single-cell contact matrices at 10 kb resolution were imputed with scHiCluster for the contacts within 5.05 Mb (result denoted as $Q_{cell}$). We only perform loop calling between 50 kb and 5 Mb, given that increasing the distance only leads to a limited increase in the number of significant loops. For each single cell, the imputed matrix of each chromosome was log-transformed, and Z-score normalized at each diagonal (result denoted as $E_{cell}$) and subtracted a local background between >=30 kb and <=50 kb (result denoted as $T_{cell}$), similar to SnapHiC (*73*). A pseudo-bulk level t-statistic was computed to quantify the deviation of E and T from 0 across single cells from the cell group, where larger deviations represent higher enrichment against global (E) or local (T) background. $E_{cell}$ is also shuffled across each diagonal to generate $E_{shufflecell}$, and then $T_{shufflecell}$, to estimate a background of the t-statistics. An empirical FDR can be derived by comparing the t-statistics of observed cells versus shuffled cells. We required the pixels to have an average E >0, fold change >1.33 against donut and bottom left backgrounds, fold change >1.2 against horizontal and vertical backgrounds (*73*), and FDR <0.01 compared to global (E) and local (T) backgrounds.

To compare the interaction strength of loops between different groups of cells, we adopt an analysis of variance (ANOVA) framework to compute the F statistics for each loop identified in at least one cell group using either $Q_{cell}$(result denoted as $F_Q$) or $T_{cell}$ (result denoted as $F_T$). Then, we Z-scored $F_Q$ and $F_T$ across all the loops being tested and selected the ones with $F_Q$ and $F_T$ > 1.036 (85th percentile of standard normal distribution) as differential loops. The threshold was decided by visually inspecting the contact maps as well as the correlation of interaction and loop anchor CG methylation.

**Identification of domains and differential domain boundaries**

Single-cell contact matrices at 25kb resolution were imputed with scHiCluster (*17*) for the contacts within 10.05 Mb. Domains were identified within each single cell. Insulation scores were computed in each cell group (major type or major type within a brain region) for each bin with the pseudo-bulk imputed matrices (average over single cells) and a window size of 10 bins. The boundary probability of a bin is defined as the proportion of cells having the bin called as a domain boundary among the total number of cells from the group.

To identify differential domain boundaries between n cell groups, we derived an nx2 contingency table for each 25kb bin, where the values in each row represent the number of cells from the group that has the bin called as a boundary or not as a boundary. We computed the Chi-square statistic and p-value of each bin and used the peaks of the statistics across the genome as differential boundaries. The peaks are defined as a local maximum of Chi-square statistics within FDR <1e-3 (Benjamini and Hochberg procedure). If two peaks are within 5 bins of each other, we only keep the peak with a higher Chi-Square statistic. We also require the peaks to have a Z-score transformed Chi-square statistic >1.960 (97.5 percentile of standard normal distribution), fold-changes between maximum and minimum insulation score >1.2, and differences between maximum and minimum boundary probability >0.05.

**Single-cell embedding with loop and domain**

We randomly selected 200 cells from each major type except for L5-ET, where only 107 cells were identified and all used in this analysis. For loops, we combined the loop pixels identified in all major types to make a meta loop list, and generated a binary cell-by-loop matrix where each element indicated whether a contact was detected in the cell at the loop pixel. Latent semantic analysis with log term frequency was applied to the binary matrix (denoted as $A$) to compute the embedding. Specifically, we selected the columns having 1 in more than 5 rows, then computed the column sum of the matrix ($colsum_j = \sum_{i=1}^{\#cell} A_{ij}$) and kept only the bins with Z-scored $log_2 colsum$ between -2 and 2. The filtered matrix was normalized by dividing the row sum of the matrix to generate a term frequency matrix $TF$, and further converted to $X$ used for singular value decomposition $X = USV^T$, where $X_{ij} = log(TF_{ij} \times 100000 + 1) \times log(1 + \frac{\#cell}{colsum_j})$. Top 15 dimensions of $U$ were then used for t-SNE visualization and cluster analysis. For domains, we generated a binary cell-by-25kb bin matrix where each element indicated whether the bin was identified as a domain boundary in the cell. The same LSI framework was used to obtain the cell embedding. K-Means was used to perform clustering, and k was enumerated from 3 to 12 and the result with the highest adjusted rand index (ARI) compared to the cluster labels was shown in Fig. 2, G and I. To benchmark the ability to separate excitatory cell types, we used L2/3-IT, L4-IT, L5-IT, L6-IT, L6-IT-Car3, L5/6-NP, L6b, L6-CT, and L5-ET. For inhibitory cell types, we used Lamp5-Lhx6, Lamp5, Sncg, Vip, Pvalb-ChC, Pvalb, and Sst.

**Integration among different single-cell datasets**

**Feature matrices for human single-cell DNA methylation, expression and open chromatin.** CG- and CH-cell type marker genes determined from the mC dataset for both major and subtype levels were used as the features for integration analysis. When integrating with scRNA (companion manuscript Siletti et al. (*14*)) or snATAC datasets (companion manuscript Li et al.), we used the opposite values of gene body methylation since they generally are strongly anti-correlated with gene expression. Both scRNA and snATAC datasets were normalized by the averaged total UMI counts of the featured genes and then transformed by *log(x+1)*. The neuronal cell types and non-neuronal cell types were integrated separately. CH-methylation was used for neuronal cell types, while CG-methylation was used for non-neuronal cell types. An additional filtering step was applied before integrating non-neuronal cell types, which required the total UMI of the featured genes of each cell to be larger than 3,000 for both scRNA and snATAC datasets.

**Feature matrices for human and mouse single-cell DNA methylation.** We used only homologous genes between human and mouse to perform the integration analysis. The list of homologous genes was downloaded from the Mouse Genome Informatics (MGI) database (http://www.informatics.jax.org/homology.shtml). The homologous genes were selected from the same features used when integrating with scRNA and snATAC datasets. Human brain cells from thalamus, midbrain, cerebellum, pons, and entorhinal cortices were excluded since no counterparts exist from the public mouse dataset (*1*). The mouse dataset was re-annotated in the same way as the human dataset. CG-methylation was used to integrate neuronal and non-neuronal cell types separately.

**Method to integrate different single-cell sequencing datasets.** After feature matrix generation, we used a 3-step method analogous to Seurat v3 to project two datasets $X$ and $Y$ onto the same space: 1) Using canonical correlation analysis (CCA) to capture the shared variance across cells between datasets; 2) finding anchors as 5 mutual nearest neighbors (MNN) between the two datasets; 3) pulling the two datasets into the same space. To allow the scalability, we randomly selected 20,000 cells from each dataset ($X_{ref}$ and $Y_{ref}$) as a reference to fit the CCA, and transform the other cells ($X_{qry}$ and $Y_{qry}$) onto the same CC space. Specifically, the canonical correlation vectors (CCV) of $X_{ref}$ and $Y_{ref}$ (denoted as $U_{ref}$ and $V_{ref}$) are computed by singular value decomposition on their dot product, $U_{ref} S V_{ref}^T = X_{ref} Y_{ref}^T$, where $U_{ref}^T U_{ref} = I$ and $V_{ref}^T V_{ref} = I$. Then the CCV of $X_{qry}$ and $Y_{qry}$ (denoted as $U_{qry}$ and $V_{qry}$) are computed by $U_{qry} = X_{qry}(Y_{ref}^T V_{ref})/S$ and $V_{qry} = Y_{qry}(X_{ref}^T U_{ref}^T)$. U and V were normalized by dividing the L2-norm of each row, and used to find MNN anchors and score anchors using the same method as Seurat v3. $X$ and $Y$ were also combined vertically and the PCs of this combined matrix were integrated together using the same method as Seurat v3 through the anchors generated from the previous step. This integration step projects the PCs of one dataset (query) to the PCs of the other dataset (reference) while keeping the PCs of the reference dataset unchanged. The resulting PCs were used for visualization and finding matched clusters between datasets.

### Enhancer prediction

Based on the pairwise CH-DMGs determined between cell subtypes, we assign a gene as hypomethylated in one subtype if it is a hypomethylated DMG in at least 40 out of 187 pairs compared with other subtypes. A DMR is assigned to a subtype if it is either CG-hypomethylated in the subtype (see section "*Determine differentially methylated regions*" above) or its CG-methylation level is below 0.3. A DMR is considered as a candidate cis-regulatory element if it is connected by a differential loop to a gene which is also a DMG in the same subtype. We do not require the differential loop connecting the DMR-DMG pair to be a loop detected in the subtype of the pair. The reason for this loose criterion is threefold: 1) the strength of the differential loop anti-correlates with the methylation levels (Fig 2K). If a DMR-DMG pair is connected with a differential loop in one subtype, the loop likely exists in another subtype with the DMR-DMG pair of similar methylation statuses; 2) Cis-regulatory elements are usually pleiotropic (*74*), The loops detected in subtypes covered by the m3c dataset could be reused in another uncovered if the methylation status of the DMR-DMG pairs is similar. 3) Loops could be missed in detection due to either the limitation of the computation methods or the insufficient coverage in certain subtypes. The DNA looping information transferring among subtypes could cope with such a situation to some extent.

### Association between brain disorder risk variants and DMRs across cell types

We obtained GWAS summary statistics for quantitative traits related to neurological disease and control traits of intelligence (*75*), educational attainment (*76*), alcohol usage(*77*), Alzheimer's Disease (*78*), bipolar disorder (*79*), attention deficit hyperactivity disorder (*80*), neuroticism (*81*), schizophrenia (*82*), amyotrophic lateral sclerosis (*83*), tobacco use disorder (*84*), insomnia (*85*), sleep duration, coronary artery disease (*86*), height, tiredness (*87*), type 1 diabetes (*88*), type 2 diabetes (*89*), allergy (*90*), birth length (*91*), and birth weight (*92*).

We prepared summary statistics to the standard format for linkage disequilibrium score regression. Next, we converted major-type hypo-DMRs to human genome assembly GRCh37 (hg19) coordinates using the software LiftOver, and annotated with the 1000 Genomes Project Phase 3 SNPs (*93*). The superset of the hypo-DMRs was used as the background. Finally, we used cell-type-specific linkage disequilibrium score regression (https://github.com/bulik/ldsc; (*38*)) to estimate the enrichment coefficient of each annotation for each trait.

### Brain regional axes from DNA methylation profiles

Both CG- and CH- highly variable 100kb-bins of one cell type (the same features for clustering analysis) were used to compute a lower dimensional representation with the principle component analysis (PCA). First, a neighbor graph of the cells was constructed in the PCA space. Then, for each cell, a regional identity vector was computed by averaging the location information of this cell and its neighbors. A pairwise Manhattan distance matrix was then constructed from the regional identity vectors to capture relations among brain regions. The principle coordinate analysis (PCoA) was applied to this distance matrix to obtain a lower-dimensional embedding in the regional space as well as preserve relative distances among cells. Thus, the cells were transformed from the methylome space to the regional space.

In the regional space, we perform the trajectory analysis with the Elastic Principal Graph (EPG) algorithm (*94*) implemented in STREAM (*46*). The parameters epg_alpha, epg_mu, and epg_lambda were manually adjusted to ensure the resulting trajectories well represented the distributions of the cells in the regional space. Each cell was assigned a regional index (or pseudotime) range in [0,1] according to its relative position to the trajectory. The cells were then grouped into 20 bins along the trajectories based on their regional index. The mean DNA methylation profiles can be computed for each bin.

**Consensus regional axis for cortex and basal ganglia.** The mean regional index was first computed for cells from each cortical region in each cell type. Then the average regional indices were calculated by averaging the mean regional indices across the corresponding cell types. Finally, the consensus regional axis was constructed by ranking the average indices.

**Regional DMGs.** We used a one-vs-rest strategy to calculate region-specific CH-DMGs (rDMGs) within major types from the cortex and basal ganglia. To avoid potential bias caused by an imbalance of cell numbers in different regions, we downsampled cells in each region to no more than 500. Using the Wilcoxon Rank Sum Test, protein-coding genes and lncRNAs were tested for significant methylation decrease (or hypomethylation). The p-values were adjusted with multitest correction using the Benjamini-Hochberg procedure. The genes with adjusted p-values$\leq 1^{-10}$ and log2 fold-change $\leq$ -0.1 were considered rDMGs.

**Regional DMRs.** Cells from the same brain region were merged for each major type to construct the regional pseudo-bulk methylation profiles. Then the DMRfind function of the software MethylPy was used to determine the candidate rDMRs with the same options as in determining cell-type DMRs. If a candidate rDMR has CG-methylation variation $\geq$ 0.6 across regions tested, it is considered an rDMR.

**Enrichment analysis on conserved DMRs between human and mouse**

**Functional enrichment analysis of hcCnsvDMRs.** The Genomic Regions Enrichment of Annotations Tool (GREAT) (*95*) was used to compute the Gene Ontology (GO) term enrichment of hcCnsvDMRS. "Basal+extension" option (5.0 kb upstream, 1.0 kb downstream, and up to 100 kb max extension) was selected for gene association, and "curated regulatory domains" are included in the analysis.

**Comparison between hcCnsvDMRs and histone modification marks in mouse forebrains.**

Replicated peaks of histone modification marks of P0 mouse forebrain were downloaded from the Encode project (*6*). Particularly, H3K27ac (ENCFF044YBD), H3K27me3 (ENCFF461UUN), H3K4me1 (ENCFF467MYU), H3K4me3 (ENCFF066LGF), and H3K9me3 (ENCFF997XJK) were used. The software Genomic Association Tester (GAT; (*96*)) was used to compute the enrichment of hcCnsvDMRs in the histone modification marks. Accessibility of hcCnsvDMRs was determined by comparing them with snATAC peaks profiled from P56 mouse brains (*2*).

**scMCode construction**

**Candidate CpG sites.** We constructed the pseudo-bulk mCG profile for each major type and then iteratively selected CpG sites to distinguish all major types except EC and PC due to the relatively low cell number of these three major types. In each iteration, CpG sites were selected according to the criteria: 1) they are either almost entirely methylated (mCG%≥80%) or unmethylated (mCG%≤20%) among all the remaining major types; 2) both two methylation statuses are presented among the remaining major types; 3) The CpG sites should have coverage ≥10 in ≥80% of the remaining major types. These selected CpG sites were added to the CpG site pool for later scMCode construction. In each iteration, the methylation levels of the selected CpGs in the remaining major types were binarized if they are ≥80% or ≤20%. Pairwise distances were computed between binarized methylation status, and the cell types that had a distance <20 to any of the other major types were kept for the next iteration of CpG selection. In total, 221,140 CpG sites were selected as candidates for scMCode construction.

**CpG site selection for scMCode.** The methylation levels of candidate CpG sites across all the major types were trinarized by if they were≥80% or ≤20%. The CpG sites were further grouped into 38,945 features based on their trinarized methylation states across major types. To prevent scMCode from bias caused by cell type population differences or individual variations of donors, we randomly select 300 cells from each major type from each donor as the dataset for scMCode construction. In each cell, the methylation state of each feature was either computed by averaging the methylation levels of all the CpG sites belonging to this feature (AverageCpG) or by directly using the methylation level of a randomly picked single CpG site belonging to this feature (RandomCpG). The cell-by-feature matrix was trinarized and then used to train a random forest (RF) model to predict major types. A 4-fold cross-validation scheme was used to prevent overfitting. Finally, the top 800 most important features were selected to construct the scMCode for major types. We observed no difference in predicting performance between AverageCpG and RandomCpG, indicating the robustness of scMCode.

**Boost the prediction accuracy with K-Nearest-Neighbor imputation.** We achieved ~88% single-cell predicting accuracy directly with the cell-by-feature matrix. Given the limited coverage of single-cell data, the cell-by-feature matrix could be further imputed to improve the prediction accuracy. Within the training dataset, we randomly selected half of the cells and merged them into pseudo-cells according to their major types. This process was repeated 20 times, and a pseudo-cell-by-feature matrix was constructed from these pseudo-cells in the same way. A K-Nearest-Neighbor (KNN) imputer was built upon this matrix. The testing dataset was first imputed with the KNN imputer when predicting cell types and then fed to the RF model. The KNN imputation step improved the prediction accuracy to ~92%.

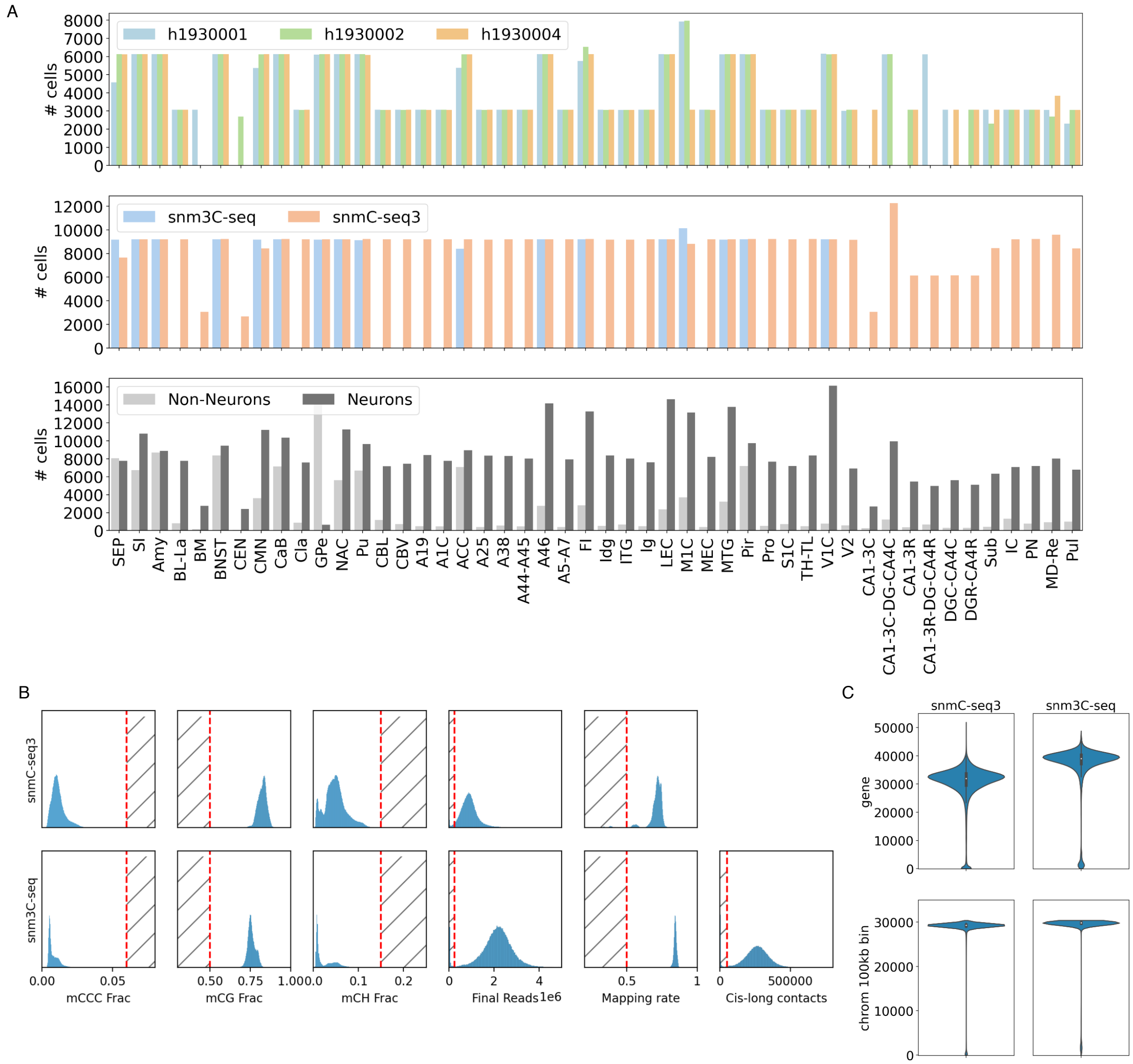


**Figure S1. Sample information and QC metrics.** (A) Cell number distributions of donors (top), epigenetic profiling assays (mid), and neuronal/non-neuronal cell types (bottom) from different brain regions. (B) QC metrics are used in filtering cells in snmC-seq and snm3C-seq. (C) Coverage per cell distributions of genomic features of genes and 100kb-bins in mC and m3C datasets.

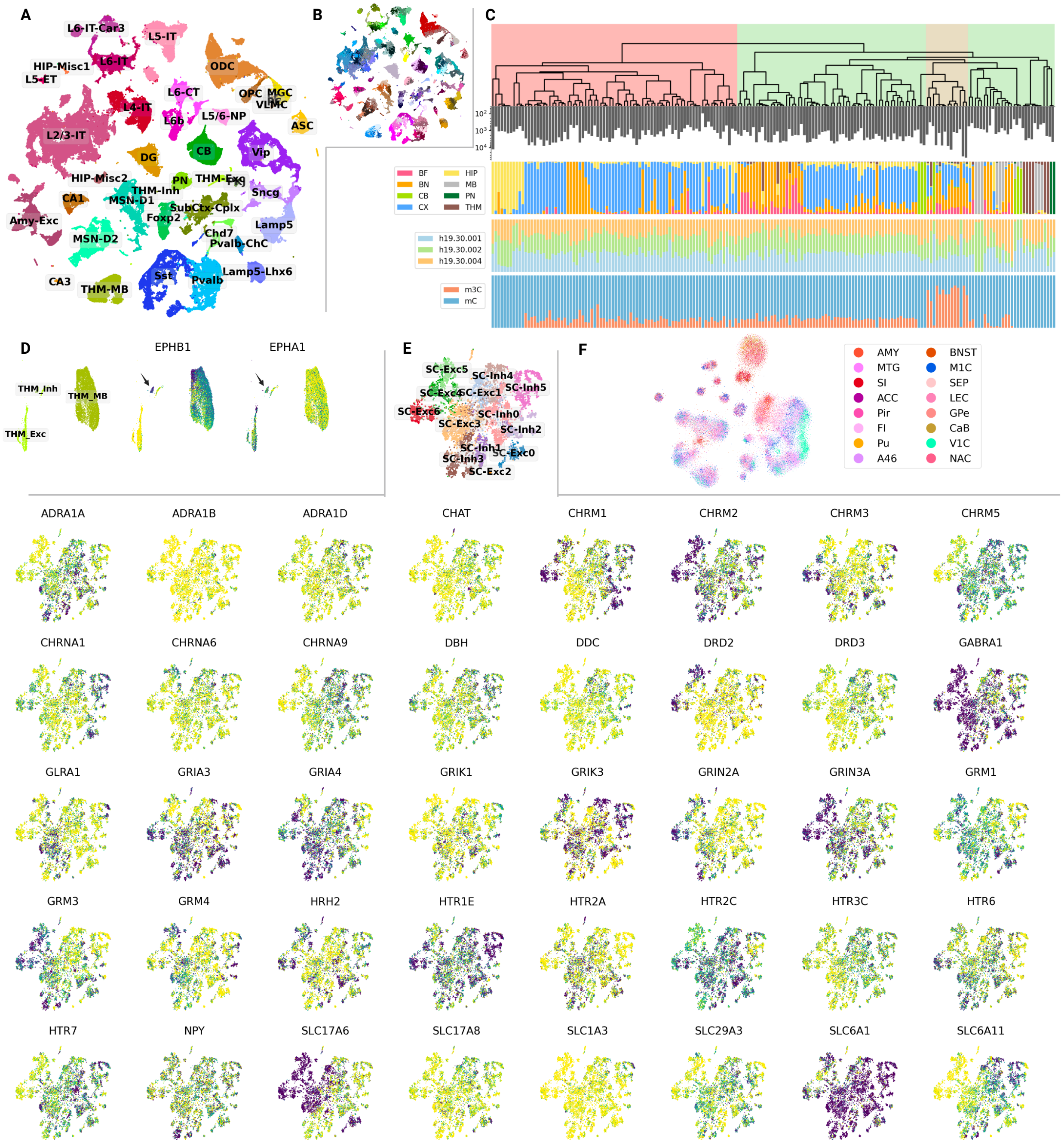


**Figure S2. Clustering and annotation of brain cell types.** (A) 2D t-SNE visualization of mC cells colored by 40 major types. (B) 2D t-SNE visualization of mC cells colored by 188 subtypes. (C) The robust dendrogram of the subtypes and the corresponding meta info of brain structures, donor origins, and epigenetic profiling assays. (D) The thalamus major type THM-Inh is hypomethylated in genes EPHB1 and EPHA1. (E) Subtypes of SubCtx-Cplx major type show highly variable DNA methylation in the genes of neurotransmitter receptors, transporters, and neuropeptides. (F) Regional diversity of single-cell embedding computed from chromatin conformation. The embedding is the same as in Fig 1G, while colored by the dissection regions.

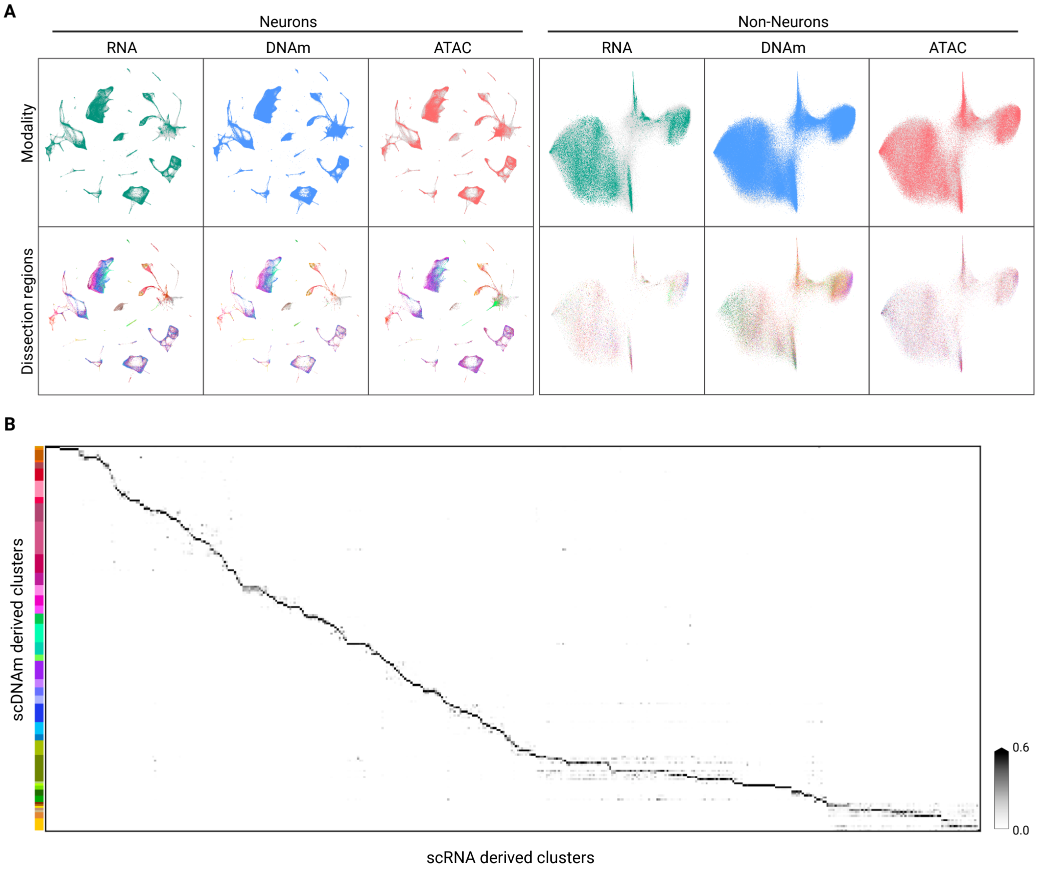


**Figure S3. Integration between modalities.** (A) 2D t-SNE visualization of integration results between snmC, scRNA, and snATAC datasets. The integration shows cell types and regional diversity are consistent among the three modalities. (B) Heatmap shows the cross-tabulation between scRNA cluster labels and transferred mC annotation. The count number was normalized by rows.

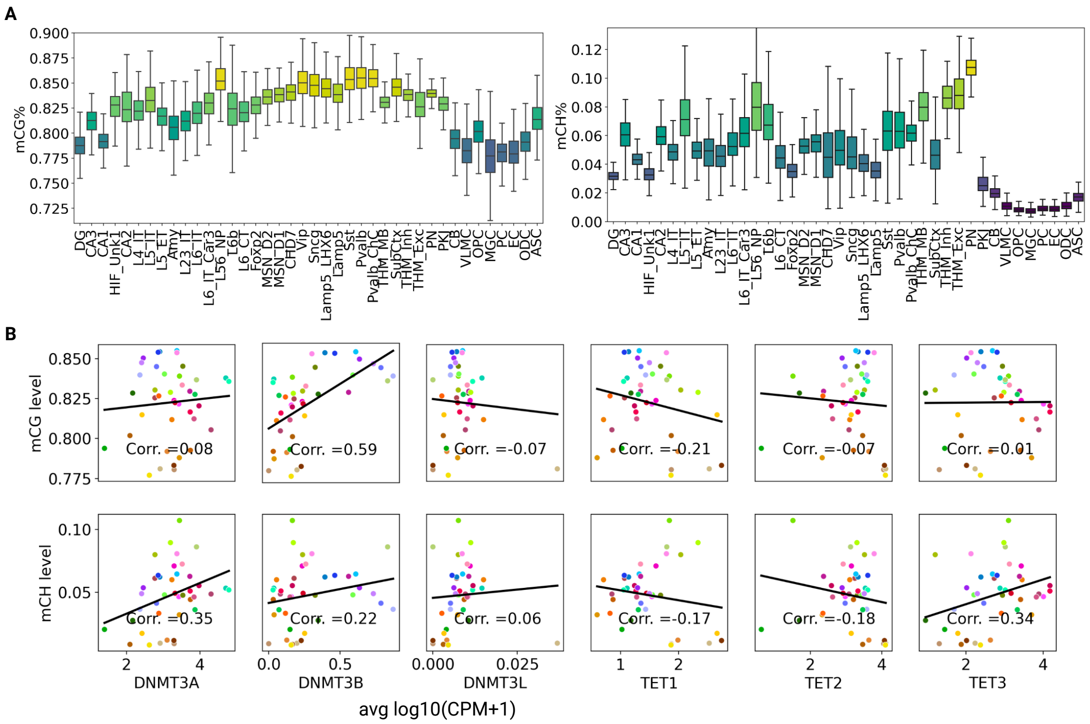


**Figure S4. Global methylation levels across major types.** (A) Global CG- and CH-methylation levels of major types. (B) Pearson correlation between global CG- and CH-methylation levels and gene expression of DNA methylation readers/modifiers across major types.

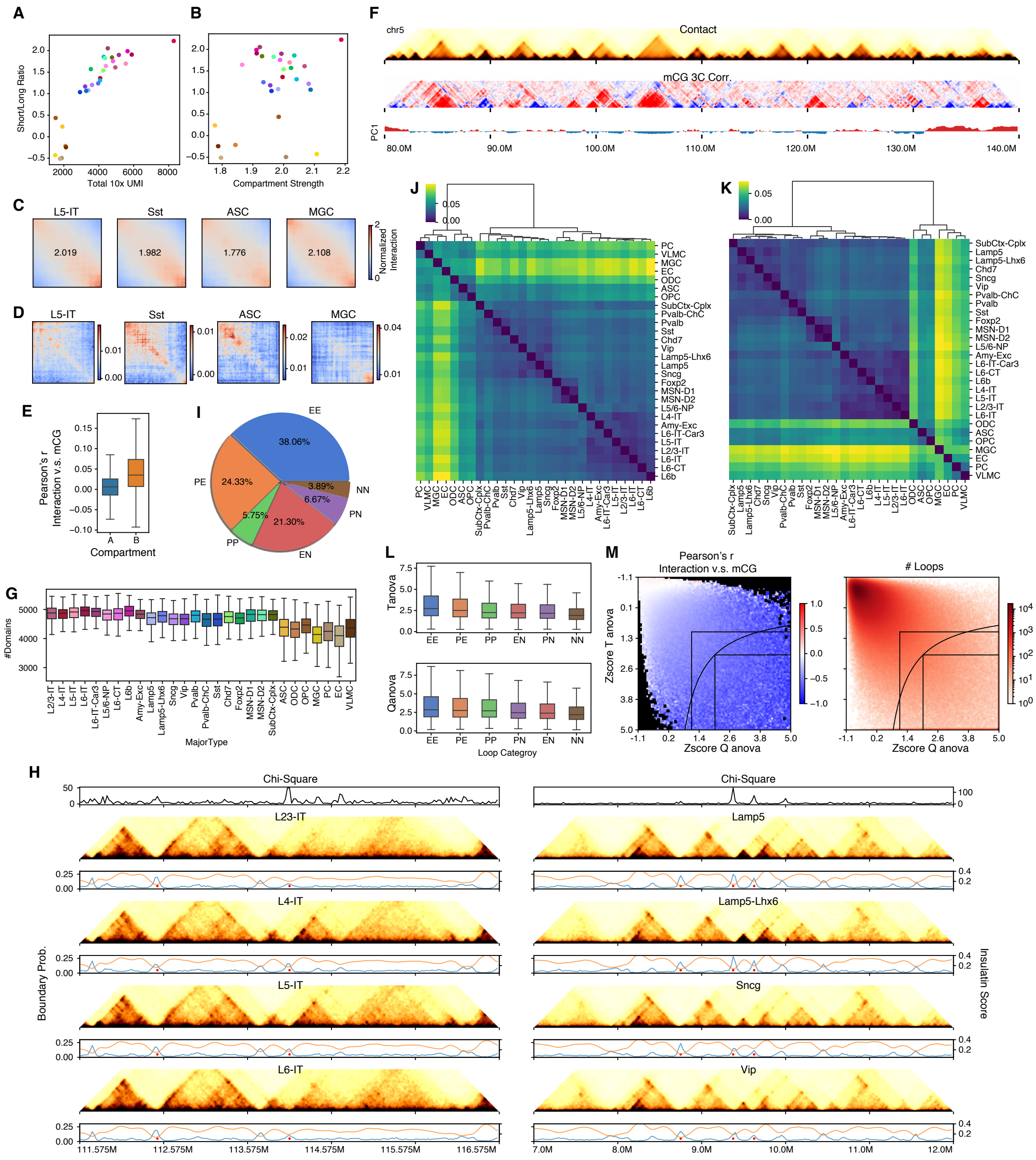


**Figure S5. Diversity of 3D genome structures across major types.** (A, B) Median log2 ratio between shorter and longer contact frequencies against average UMI detected per cell (A) or compartment strength computed from the imputed matrix for each cell type (B). The color palettes are shared across the whole manuscript. (C, D) Saddle plots of four cell types are shown in Fig. 2C to 2E. The axes are ranked by the compartment score of the cell types. Values are distance-normalized contacts (C) or mCG correlation (D). (E) Imputed contact map (top), the correlation between 3C and mCG (middle), compartment score computed from raw contact matrices (bottom) at 100kb resolution. (F) Correlation between 3C and mCG in the triangle region of A compartment segments (n=346,260) and B compartment segments (n=172,045). (G) Number of domains identified in single cells of each major type. (H) The imputed contact matrices (heatmap), boundary probabilities (blue lines), insulation scores (orange lines), and differential boundaries (red dots) are centered at the TSS of FOXP2 (left) or LAMP5 (right). (I) Proportion of different categories of loops. P denotes loop anchors overlapping with promoters (TSS±2k), E denotes loop anchors overlapping with DMRs but not promoters, and N denotes loop anchors overlapping with neither promoters nor DMRs. (J, K) Cosine distances between major types are measured by imputed contact strengths across all loop pixels identified in at least one cell type (J) or boundary probabilities across all 25kb bins (K). (L) ANOVA statistics of different categories of loop pixels are computed with T (top) or Q (bottom). (M) The correlation between interaction strength and average mCG level at two anchors (left) or the number of loop pixels (right) with different T and Q ANOVA statistics.

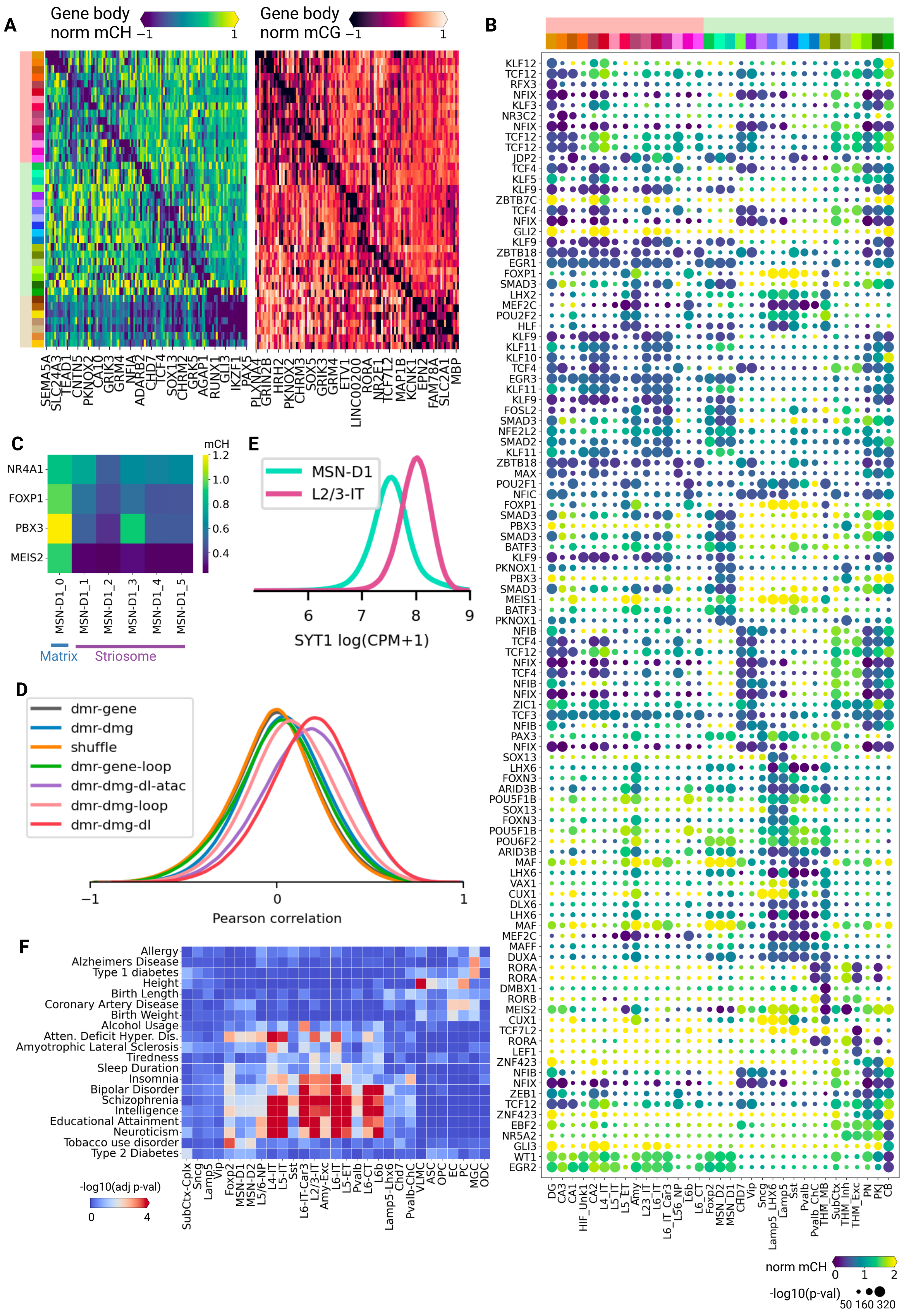
A
Gene body norm mCH −1 1
Gene body norm mCG −1 1
SEMA5A SLC24A3 TEAD1 CNTN5 PKNOX2 CA10 GRIK3 GRM4 NFIA ADARB2 CHD7 TCF4 SOX13 CHRM3 GRK5 AGAP1 RUNX1 GLI3 IKZF1 PAX5
PLXNA4 GRIN2B HRH2 PKNOX2 CHRM3 SOX5 GRIK3 GRM4 ETV1 LINC00200 RORA NR2F1 TCF7L2 MAP1B KCNK1 EPN2 FAM78A SLC2A1 MBP
B
KLF12 TCF12 RFX3 NFIX KLF3 NR3C2 NFIX TCF12 TCF12 JDP2 TCF4 KLF5 KLF9 ZBTB7C TCF4 NFIX GLI2 KLF9 ZBTB18 EGR1 FOXP1 SMAD3 LHX2 MEF2C POU2F2 HLF KLF9 KLF11 KLF10 TCF4 EGR3 KLF11 KLF9 FOSL2 SMAD3 NFE2L2 SMAD2 KLF11 ZBTB18 MAX POU2F1 NFIC FOXP1 SMAD3 PBX3 SMAD3 BATF3 KLF9 PKNOX1 PBX3 SMAD3 MEIS1 BATF3 PKNOX1 NFIB TCF4 TCF12 NFIX TCF4 NFIB NFIX ZIC1 TCF3 NFIB PAX3 NFIX SOX13 LHX6 FOXN3 ARID3B POU5F1B SOX13 FOXN3 POU5F1B POU6F2 ARID3B MAF LHX6 VAX1 CUX1 DLX6 LHX6 MAF MEF2C MAFF DUXA RORA RORA DMBX1 RORB MEIS2 CUX1 TCF7L2 RORA LEF1 ZNF423 NFIB NFIX ZEB1 TCF12 ZNF423 EBF2 NR5A2 GLI3 WT1 EGR2
DG CA3 CA1 HIF_Unk1 CA2 L4_IT L5_IT L5_ET Amy L23_IT L6_IT L6_IT_Car3 L56_NP L6b L6_CT Foxp2 MSN_D2 MSN_D1 CHD7 Vip Sncg Lamp5_LHX6 Lamp5 Sst Pvalb Pvalb_ChC THM_MB SubCtx THM_Inh THM_Exc PN PKJ CB
norm mCH 0 1 2
-log10(p-val) 50 160 320
C
mCH
NR4A1 FOXP1 PBX3 MEIS2
MSN-D1_0 MSN-D1_1 MSN-D1_2 MSN-D1_3 MSN-D1_4 MSN-D1_5
Matrix Striosome
1.2 1.0 0.8 0.6 0.4
E
MSN-D1
L2/3-IT
6 7 8 9
SYT1 log(CPM+1)
D
dmr-gene
dmr-dmg
shuffle
dmr-gene-loop
dmr-dmg-dl-atac
dmr-dmg-loop
dmr-dmg-dl
−1 0 1
Pearson correlation
F
Allergy
Alzheimers Disease
Type 1 diabetes
Height
Birth Length
Coronary Artery Disease
Birth Weight
Alcohol Usage
Atten. Deficit Hyper. Dis.
Amyotrophic Lateral Sclerosis
Tiredness
Sleep Duration
Insomnia
Bipolar Disorder
Schizophrenia
Intelligence
Educational Attainment
Neuroticism
Tobacco use disorder
Type 2 Diabetes
SubCtx-Cplx Sncg Lamp5 Vip Foxp2 MSN-D1 MSN-D2 L5/6-NP L4-IT L5-IT Sst L6-IT-Car3 L2/3-IT Amy-Exc L6-IT L5-ET Pvalb L6-CT L6b Lamp5-Lhx6 Chd7 Pvalb-ChC VLMC ASC OPC EC PC MGC ODC
-log10(adj p-val)
0 4

**Figure S6. Gene regulation in brain cells.** (A) Different major types have specific marker genes in both CG- and CH-methylation. All marker genes shown in the heatmaps are TFs, neurotransmitter receptors, transporters or neuropeptides.(B) The scatter plot of CH-methylation and enrichment of TFs that were assigned to the major types. (C) Heatmap shows average CH-methylation levels of striosome markers among MSN-D1 subtypes. The subtypes MSN-D1 1-5 are hypomethylated in these genes, indicating they are likely from the striosome compartment of striatum. (D) Distribution of Pearson correlations between CG-methylation levels of DMRs and CH-methylation levels of genes. Consideration of differentiation of gene body methylation and DNA loops greatly improves the association between DMRs and genes. (E) Distribution of SYT1 expressions in MSN-D1 and L2/3-IT. L2/3-IT cells have high expression levels. (F) Heatmap showing the results of linkage disequilibrium score regression analysis of the variants associated with the indicated traits or diseases in loop-overlapped DMRs identified from human major cell types.

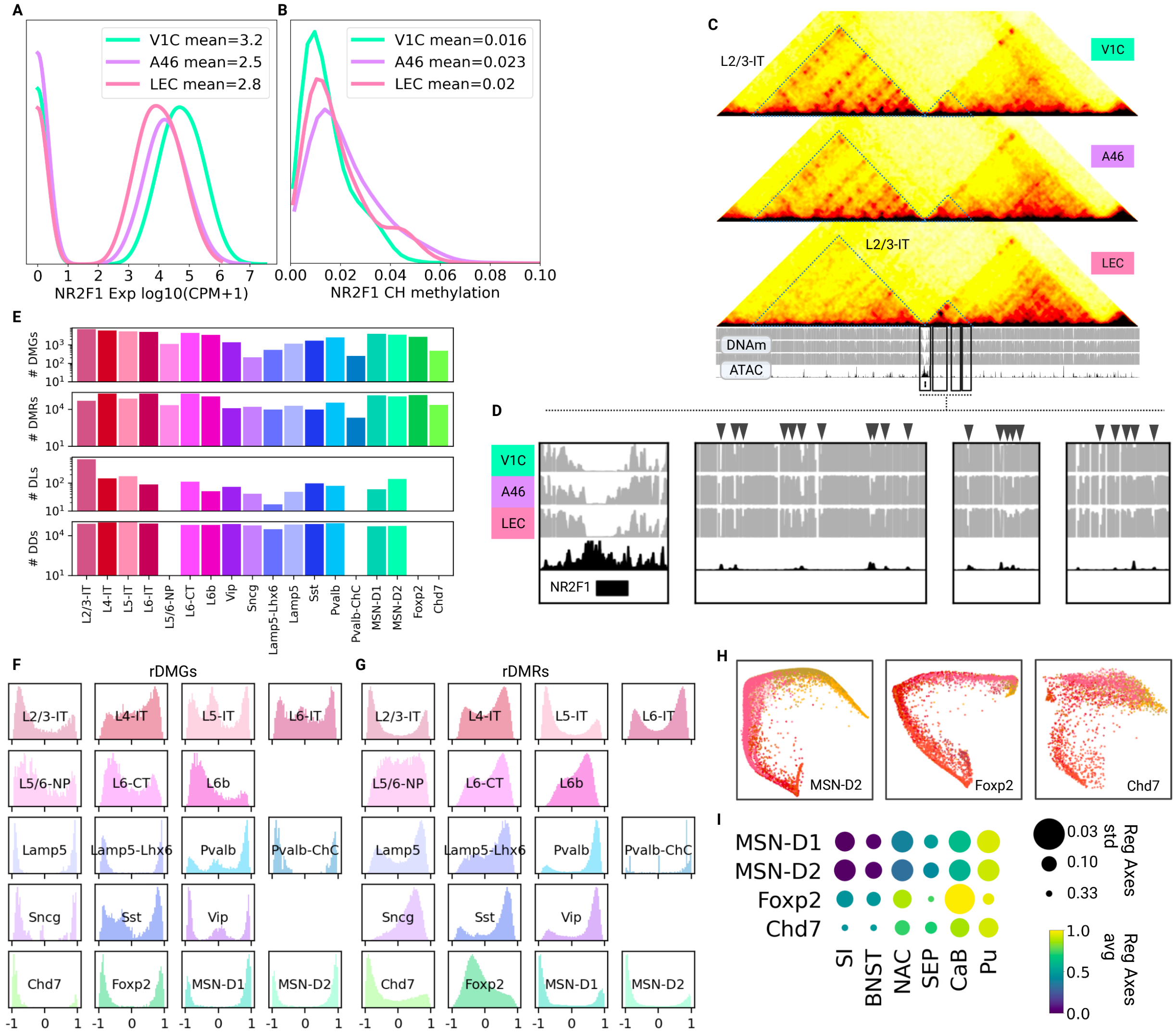


**Figure S7. Regional axes of cortical and subcortical cells.** (A) The gene NR2F1 has higher expression levels in L2/3-IT cells from V1C and LEC than A46. (B) It also has concordant lower CH-methylation levels in L2/3-IT cells from V1C and LEC than A46. (C) Chromatin conformation around the gene NR2F1 shows gradient changes in domain and loop strength. The two associated chromatin domains change in opposite directions. (D) Zoom-in view of example differential-loop-overlapping rDMRs from the "increasing" domain. The methylation levels decrease from V1C to A46 to LEC. (E) The number of regionally differential features is shown in barplots. (F, G) Distributions of Pearson correlations between methylation levels of rDMG (CH, left) or rDMR(CG, right) and regional axes determined in cortical regions (left) and basal ganglia (right), respectively. Considerable features show methylation gradients along the axes, manifested by Pearson correlations close to -1 or 1. (H) The embedding of major types MSN-D2, Foxp2, and Chd7 in regional spaces colored by dissection regions show that they share similar regional axes. (I) The consensus regional axis of basal ganglia constructed in the same way as in Fig 4C.

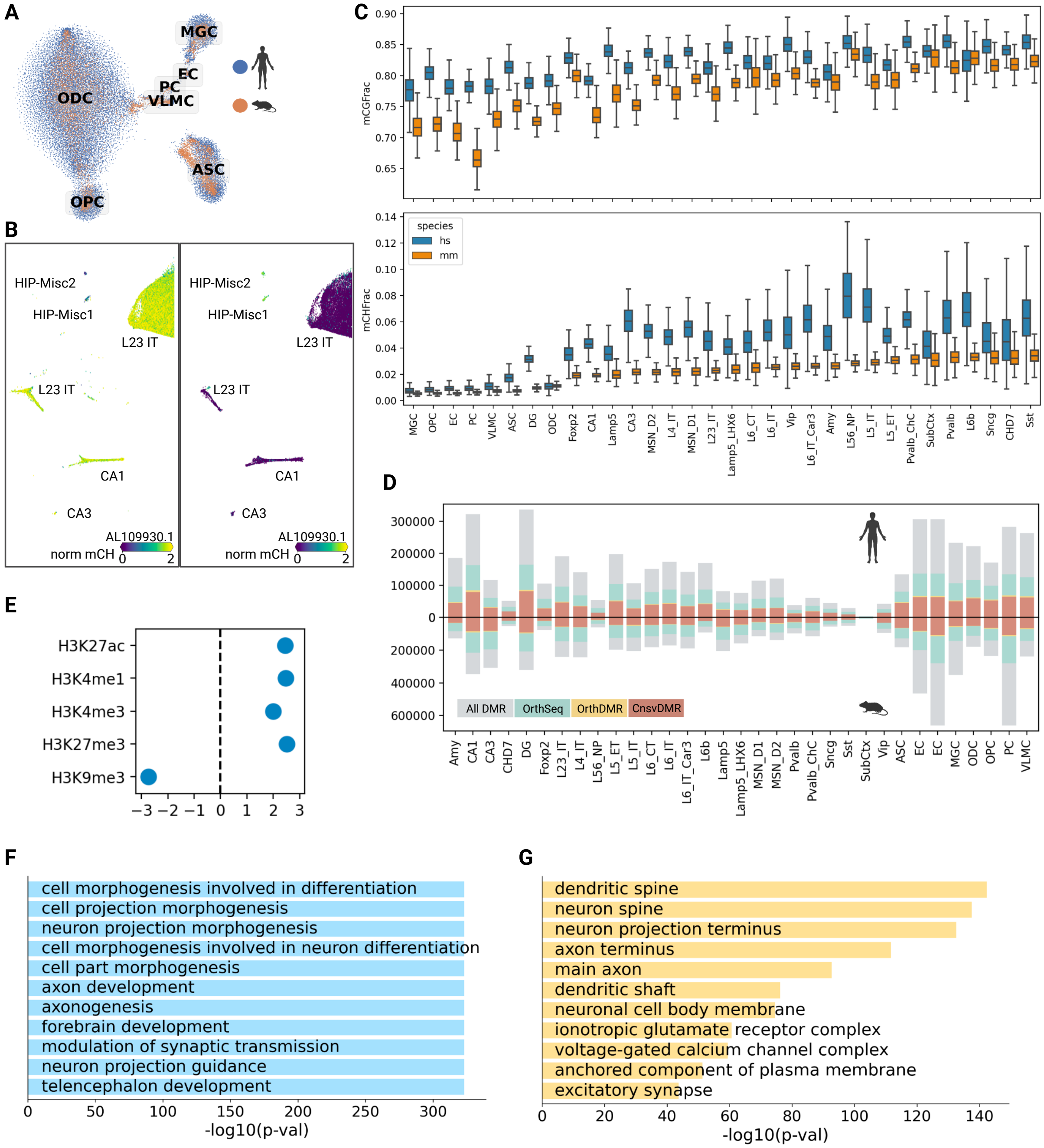


**Figure S8. Cross-species comparison between human and mouse brain cell methylomes.** (A) Integration of single-cell methylomes between human and mouse brains shows cell type conservation across species in non-neurons. (B) The cell types of HIP-Misc1 and HIP-Misc2 both feature CH-hypomethylation and gene expression of lncRNA AL109930.1. (C) Boxplots show a detailed comparison of global CG- and CH-methylation levels of conserved cell types between the human and mouse. (D) Cell type-specific numbers of DMRs in different cross-species matching categories. (E) Comparison to histone modification marks in mouse forebrains shows that the hcCnsvDMRs are depleted from heterochromatic regions (H3K9me3) as well as enriched in regions of enhancers (H3K27ac & H3K4me1), promoters (H3K4me3), and poised enhancers (H3K27m3) (F, G) GO term enrichment analysis show that hcCnsvDMRs are highly enriched in biological processes related to forebrain development (F) and in cellular components related to dendrites and synapses (G).

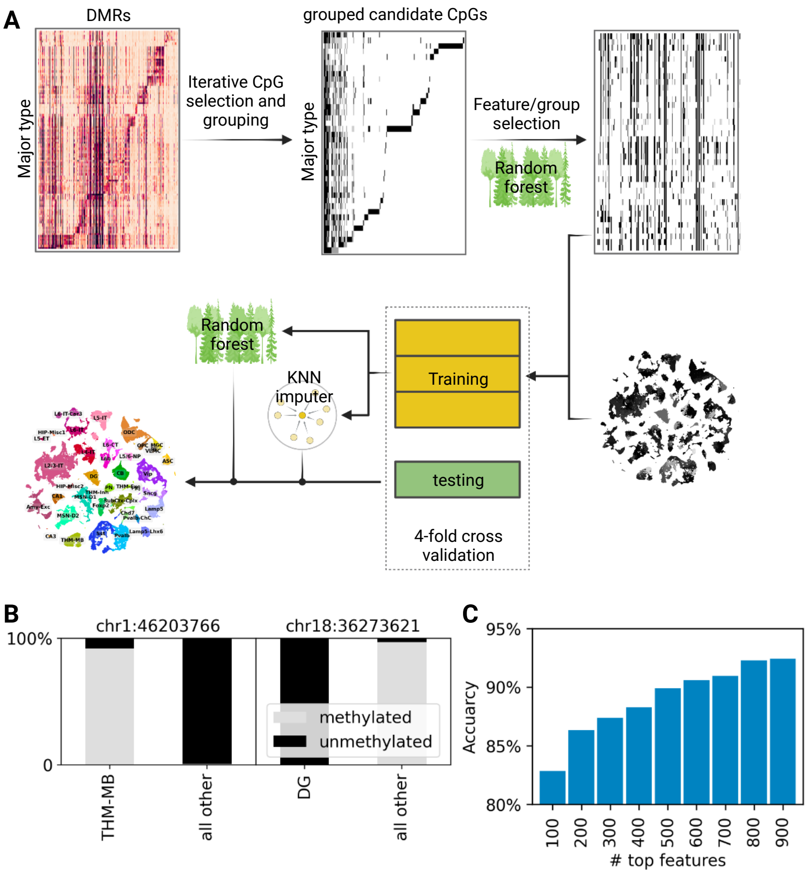


**Figure S9. snMCodes for brain cell types.** (A) Detailed workflow of the derivation of snMCodes (Methods). (B) Example of highly cell-type-specific differentially methylated CpG sites. (C) Prediction accuracy of snMCodes increases with the number of features used, which show saturation around 800~900.

**Table S1. Information of samples**

**Table S2. Brain regions and abbreviation**

**Table S3. Information of brain donors**

**Table S4. Cell class, major types and subtypes**

**Table S5. Cell meta information and annotation of single nuclei profiled with snmC-seq3**

**Table S6. Cell meta information and annotation of single nuclei profiled with snm3C-seq**

**Table S7. CpG sites and groups of scMCodes**

**Table S8. Methylation status of feature groups of scMCodes**
